# TIP-seq: A Single-cell Multiomics Approach for Simultaneous Transcriptome and Intracellular Protein Profiling

**DOI:** 10.1101/2025.05.26.653374

**Authors:** Yingying Xue, Jingyu Qi, Guangting Xie, Mingsheng Wang, Wenhua Xu, Chaoqun Ma, Xueqi Xu, Yuan Yang, Ying He, Yu Guo, Shaoyi Chen, Zhiyin Chen, Mantang Qiu, Changjiang Feng, Jianfeng Liu, Hao Wu, Ya Liu, Tao Xu, Xiaobao Cao

**Affiliations:** College of Life Science and Technology, Huazhong University of Science and Technology, Wuhan, 430074, China; Guangzhou National Laboratory, Guangzhou,510005, China; College of Life Sciences, University of Chinese Academy of Sciences, Beijing, 101408, China; School of Mechanical & Automotive Engineering, South China University of Technology, Guangzhou, 510641, China; School of Life Sciences, Nankai University, Tianjin, 300071, China; Lead Healthcare (Guangzhou) Co., LTD, Guangzhou, 510005, China; Cancer Hospital & Shenzhen Hospital, Chinese Academy of Medical Sciences and Peking Union Medical College, Shenzhen, 518116, China; Department of Thoracic Surgery, Thoracic Oncology Institute, Peking University People’s Hospital, Beijing, 100044, China; Department of Thoracic Surgery, Shenzhen Second People’s Hospital, Shenzhen, 518035, China; Department of Thoracic Surgery, The First Medical Center of Chinese PLA General Hospital, Beijing, 100853, China

**Keywords:** Single-cell multiomics, Transcriptomics, Intracellular proteins, Cell heterogeneity, Tumor microenvironment

## Abstract

Single-cell multiomics technologies have significantly advanced our understanding of cellular heterogeneity; however, the concurrent profiling of mRNA transcripts and target proteins continues to pose a substantial challenge. To bridge the gap between single-cell sequencing and proteomic methods, we developed an innovative single-cell multiomics methodology, termed TIP-seq, which facilitates the simultaneous profiling of the transcriptome and intracellular proteins without prior cell manipulation. In TIP-seq, microfluidic technology is leveraged to integrate oligonucleotide-labeled antibodies with hydrogel beads, allowing for the capture of mRNAs and proteins from individual cells within droplets, thereby enabling high-throughput and precise dual-omics data acquisition. When applied to lung cell lines and non-small cell lung cancer (NSCLC) tissues, TIP-seq analysis revealed notable cellular heterogeneity and molecular dynamics, emphasizing the distinct immune cell interactions within NSCLC tissues. Key immune checkpoint interactions in NSCLC, such as SPP1–CD44, NECTIN2–TIGIT, and NECTIN2–CTLA4, along with functional alterations in tumor-associated dendritic cells (DCs) and T cells, were identified via TIP-seq, underscoring their pivotal roles in mediating immune suppression within the tumor microenvironment. Collectively, TIP-seq represents a powerful methodology for identifying novel therapeutic targets and biomarkers, thereby holding significant potential for the advancement of precision medicine in the treatment of lung cancer and other complex diseases.

## Introduction

Cellular heterogeneity and the dynamic tumor microenvironment (TME) constitute fundamental drivers of cancer progression and therapeutic resistance in solid tumors^1–3^. Although single-cell transcriptomics has significantly revolutionized our understanding of the TME and its characteristics^4, 5^, its inability to quantify intracellular proteins leaves genotype-phenotype relationships unelucidated^6, 7^. This gap is particularly critical in the field of cancer immunology, where posttranslational modifications, such as phosphorylation, often dictate protein localization and function independently of mRNA expression levels^8–10^. Such discrepancies frequently lead to misguided therapeutic targeting^6, 11, 12^, underscoring the urgent need for simultaneous transcriptome-proteome profiling to accurately capture true functional states.

Single-cell RNA sequencing (scRNA-seq)^13–15^ has been instrumental in elucidating cellular heterogeneity and identifying novel cell types and states. This technique offers a comprehensive overview of the transcriptome, allowing for the identification of differentially expressed genes and regulatory networks. Nonetheless, scRNA-seq is limited by its inability to collect protein expression data, which is essential for understanding functional cellular states. Conversely, single-cell mass cytometry (CyTOF)^16–18^ allows the simultaneous detection of several tens of proteins at single-cell resolution but is not designed to collect transcriptomic data.

In recent years, the rapid evolution of single-cell multiomics technologies has profoundly enhanced our understanding of molecular heterogeneity at the cellular level. Similar to CITE-seq^7^ and REAP-seq^19^, notable advancements have emerged by integrating scRNA-seq with antibody-oligonucleotide conjugates (AOCs), thereby allowing the simultaneous profiling of transcriptomes and surface proteins^20–22^. In secretion encoded single-cell sequencing (SEC-seq)^23, 24^, the secretions of single cells captured in hydrogel nanovials are stained with fluorescently labled surface-modified antibodies. This approach integrates microfluidics and scRNA-seq to facilitate high-throughput coanalysis of transcriptomes and secreted proteins at the single-cell level. Similarly, time-resolved assessment of protein secretion from single cells by sequencing (TRAP-seq)^25^ utilizes antibody dimers to anchor capture antibodies to the cell surface. Single cells are isolated into microwells and incubated with the antibodies to stain the secreted proteins, followed by multiomics profiling. However, owing to the fluidity of the cell membrane, the capture efficiency of TRAP-seq is contingent upon the ability of the antibody dimers to robustly bind membrane proteins, which directly impacts sensitivity. Other methodologies, such as antibody array-based chips in large-scale arrays, provide comprehensive insights into cellular states. Nevertheless, both approaches are limited by their inability to profile intracellular proteins and require prior cell manipulation, which may introduce variability and limit their applicability to a narrow range of clinical samples^26, 27^.

Multiplexed protein and transcriptomic technologies^28^, such as those combining proximity extension assays with reverse transcription, enable the simultaneous analysis of transcriptomes and intracellular proteins at the single-cell level. However, these approaches necessitate the preisolation of cells into microwells for lysis, which complicates experimental workflows and may introduce biases. Additionally, their throughput is constrained, rendering them unsuitable for large-scale studies. Techniques such as single-cell RNA and Immunodetection (RAID)^29^ and intracellular staining and sequencing (INs-seq)^30^ employ AOCs and reversible fixation for combined transcriptome and intracellular protein analysis but require intricate optimization of fixation and staining procedures.

In this study, we introduce an innovative platform termed TIP-seq, which facilitates the concurrent analysis of **t**ranscriptomics and **i**ntracellular **p**roteins through **seq**uencing (**TIP-seq**). This platform enables the profiling of individual cells without the need for labeling or fixation. This innovative approach leverages microfluidic technology to encapsulate cells, antibody-coated beads, and AOCs within droplets, thus facilitating the simultaneous tagging of RNA and intracellular proteins from a single cell. By integrating these two layers of molecular information, TIP-seq provides a comprehensive perspective on cellular function and heterogeneity. We validated the platform using A549 cell lines and non-small cell lung cancer (NSCLC) tissues, demonstrating its ability to reveal discrepancies between mRNA and protein expression. TIP-seq overcomes the limitations of existing single-cell multiomics approaches and offers new insights into disease mechanisms and potential therapeutic targets, thereby paving the way for more precise and personalized medical interventions.

## Results

### Development of TIP-seq: A Novel Single-cell Multiomics Platform

Intracellular proteins are crucial for cellular function, serving as signaling molecules that drive cell proliferation and growth, as well as regulating gene expression to influence cell differentiation. Variations in their levels can serve as biomarkers for disease diagnosis, treatment monitoring, and prognosis assessment. In response, we have developed an innovative technology that enables the simultaneous analysis of intracellular proteins and the transcriptome at the single-cell level, providing a powerful tool and novel perspective for investigating disease-associated proteins and their gene expression profiles.

To label the transcripts and proteins from individual cells, microfluidic technology was used to encapsulate antibody-coated beads, cells, AOCs, and hydrogel beads within droplets. Upon cell lysis within the droplet, target proteins and mRNA transcripts are released, while the hydrogel bead simultaneously dissolves to liberate the barcoding oligonucleotides that contain two distinct capture sequences (**Figure 1d**). The mRNAs subsequently anneals to the complementary poly-T barcoding oligonucleotides and undergoes reverse transcription within the droplets for 2 hours at 42℃ (**Figure 1b**). To isolate the target proteins, we adopt an immuno-PCR approach that involves the use of antibody-coated beads to capture the proteins of interest. Therefore, in addition to capturing intracellular proteins TIP-seq can also effectively isolate membrane proteins. These proteins are then tagged with AOCs, which imparts a unique barcode to the protein. In addition, these AOCs are tagged with additional barcoding oligonucleotides (**Figure 1b**). This process ensures that both the target proteins and mRNAs from a single cell are marked with a consistent cellular identity marker. Upon disruption of the emulsion, we magnetically isolated the protein information and recovered the reverse-transcribed products from the supernatant (**Figure 1c**). The subsequent construction of a PCR library and sequencing facilitate the acquisition of single-cell-level data pertaining to both the target proteins and the transcriptome (**Figure 1e**).

**Figure 1:**
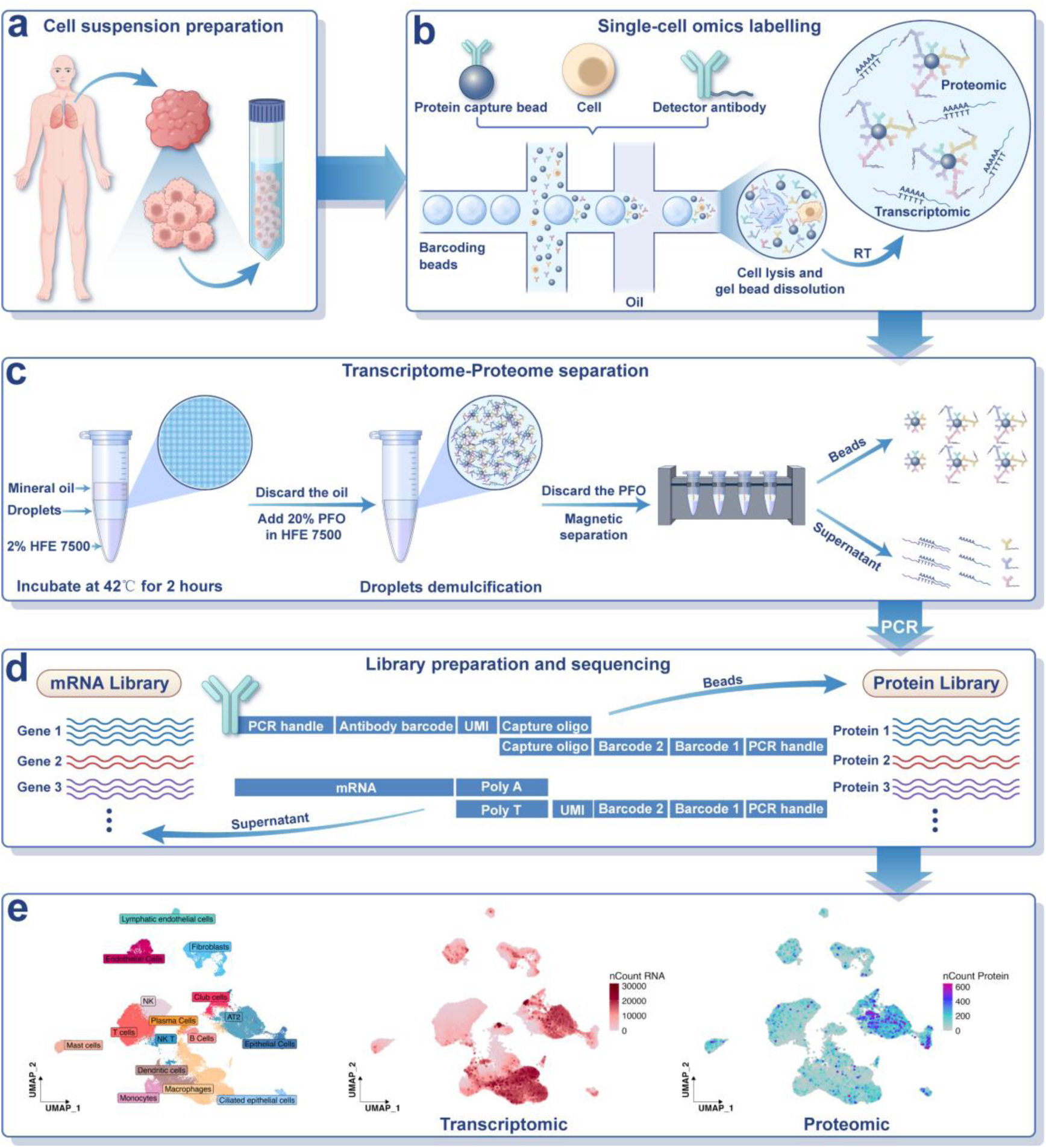
Workflow of TIP-seq. (a) Tissue dissociation to generate single-cell suspensions. (b) Barcoding of single-cell transcriptomes and proteins within droplets. (c) Droplet demulsification for splitting transcriptomes and proteins. (d) Design of barcoding oligonucleotides. (e) Distribution patterns of mRNA and protein expression.

### Validation of TIP-seq in Cell Lines

To prepare barcoding hydrogel beads, a three-dimensional network that connects linear acrylamide polymers was formed using bis(acrylamide)-L-cysteine (BAC) as a crosslinker. Additionally, acrylamide-modified oligonucleotides were incorporated into droplets formed during the mixture of BAC and acrylamide polymers. These droplets were subsequently solidified at 65℃, followed by demulsification and washing to produce hydrogel beads containing PCR handles (**Figure 2a**). By utilizing a split-and-pool strategy, we attached two barcode segments to these beads. Upon exposure to reducing agents such as THPP, the disulfide bonds within BAC were cleaved, resulting in the dissolution of the beads and the release of the barcoding oligonucleotides. As illustrated in **Figure 2b**, the hydrogel beads exhibited a uniform size and dispersion, with diameters concentrated in the range of 50–54 µm (**Figure 2c**), which are essential for ensuring consistent performance. To increase the encapsulation efficiency of the hydrogel beads, we optimized the parameters of the microfluidic chip and the injection pressure, achieving an encapsulation rate of 90% (**Figure 2d**). With the co-analysis platform established, we validated its feasibility using the 293T and 3T3 cell lines. The cross-contamination rate among these cell lines was determined to be less than 3% (**Figure 2e**), conffirming the reliability of our experimental results. Furthermore, the uniform manifold approximation and projection (UMAP) clustering of human peripheral blood cells appeared normal (**Figure 2f**), indicating the applicability of the platform across various cellular contexts. Additionally, a comparative analysis of TIP-seq and MobiNova revealed that both technologies identified a comparable number of cell types and revealed similar cell clustering results, with a high correlation coefficient of 0.93 (**Figure S3**). To further evaluate the robustness of our platform, we performed dual-omics information recovery experiments using the A549 cell line (**Figure 2g**). The proteomic data acquired were corroborated through a bulk enzyme-linked immunosorbent assay (ELISA) (**Figure 2h**, **Figure 2i**, **Figure 2j**), confirming the validity of TIP-seq protein measurements.

**Figure 2:**
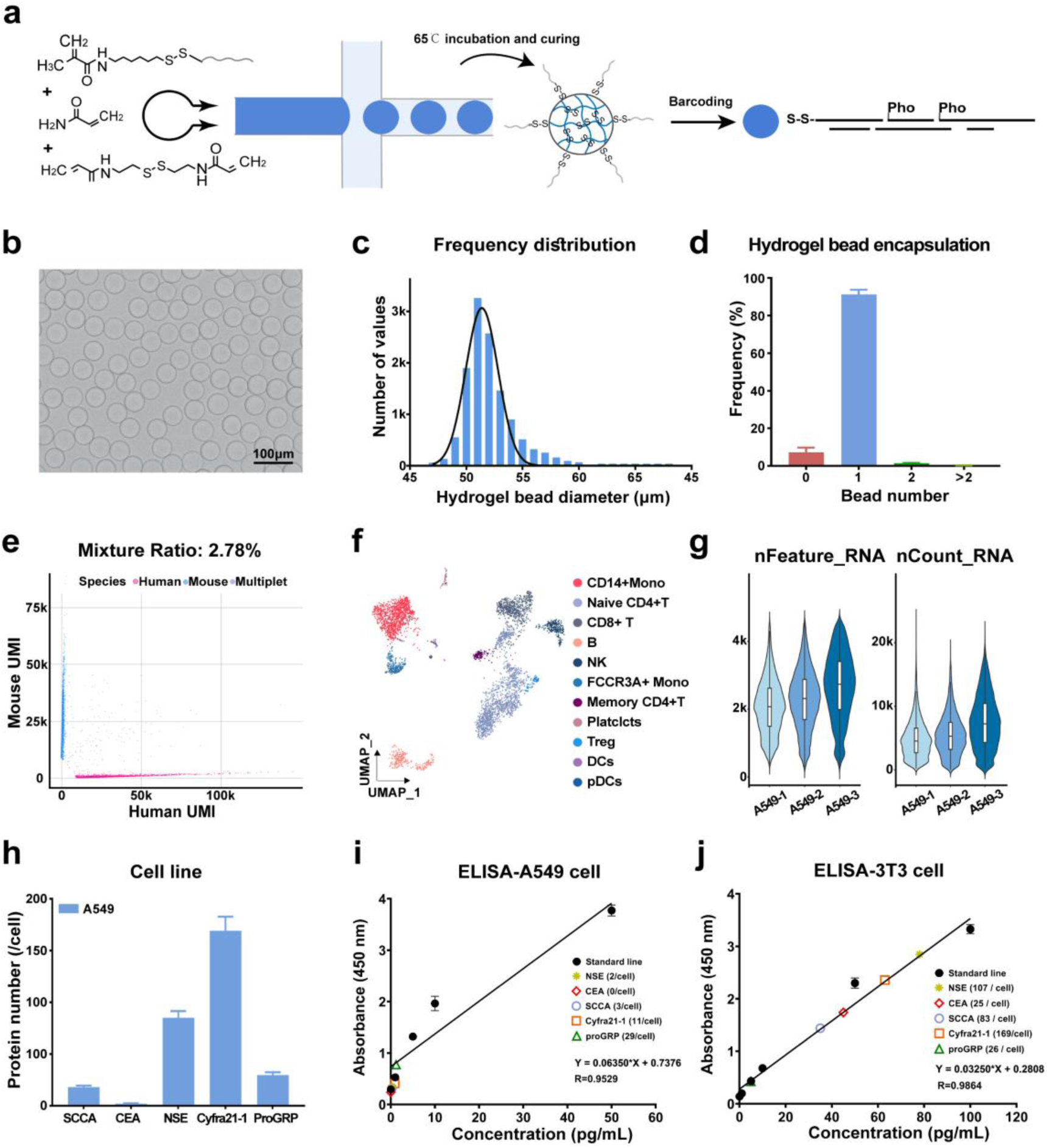
Characterization and application of hydrogel beads in TIP-seq. (a) Schematic depiction of the generation and barcoding process for hydrogel beads. (b) Representative image illustrating the morphology of the hydrogel beads. (c) Histogram showing the size distribution of the hydrogel beads. (d) Analysis of the encapsulation efficiency of hydrogel beads within droplets. (e) TIP-seq analysis of human and mouse cell mixtures. (f) UMAP visualization of human PBMC clustering on the basis of TIP-seq data. (g) Violin plots showing the distribution of gene counts and unique molecular identifiers (UMIs) per cell across three independent biological replicates (A549-1, A549-2, A549-3) in the A549 cell line. (h) Quantification of lung cancer protein biomarkers in the A549 cell line using TIP-seq, presented as a bar plot indicating the average protein content per cell. ELISA analysis of protein biomarkers associated with lung cancer in cellular extracts (i) A549 cell line. (j) 3T3 cell line.

### Application of TIP-seq in NSCLC: Protein Tumor Marker and Transcriptome Analyses

NSCLC is the predominant subtype of lung cancer and is characterized by aggressive metastasis and notable drug resistance, both of which significantly contribute to unfavorable clinical outcomes. In the present study, we applied TIP-seq technology to elucidate the molecular landscape of NSCLC by collecting paired tumor and adjacent normal tissues from four lung adenocarcinoma (LUAD) patients (**Table S2**, patients 1-4) to evaluate their transcriptomes and the expression of five specific protein tumor markers at single-cell resolution. As illustrated in **Figure S4**, the intragroup correlation coefficients among samples from all four patients exceeded 0.93, highlighting the robustness and reproducibility of our experimental methodology. These finding reinforce the validity of our findings and establish a foundational framework for further investigations into the biology of NSCLC and the identification of potential therapeutic targets.

We delineated the distribution of cellular populations within tissue samples (**Figure 3a**) and assessed the expression patterns of key marker genes (**Figure S5b**). These findings were further substantiated through dot plots and heatmaps, which validated the annotations of the cell types (**Figure S5c-d**). The gene expression profiles and protein expression of tumor markers in NSCLC tissue samples are depicted in **Figures 3b** and **3c**, respectively. For these tumor biomarkers, significant discrepancies were observed between the mRNA expression levels and the protein expression levels. Specifically, although the transcriptome data indicated markedly high expression of these tumor biomarkers in both macrophages and alveolar type 2 (AT2) cells, protein tumor markers was predominantly expressed within the AT2 cell population. These findings prompted a comprehensive investigation into the molecular and functional characteristics of these two cell types.

**Figure 3.**
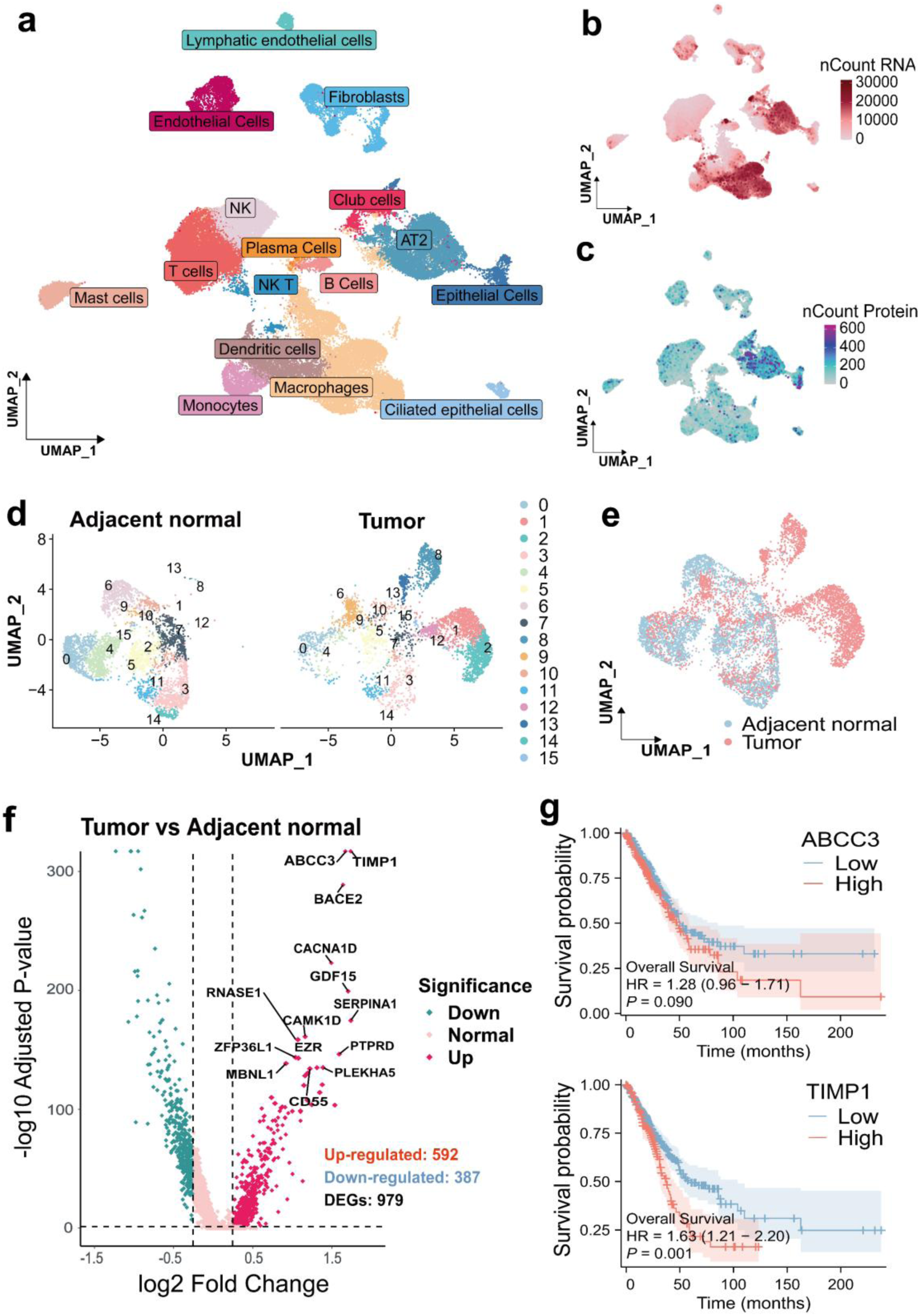
TIP-seq analysis of protein tumor markers and transcriptomes in NSCLC. (a) UMAP visualization illustrates the cluster assignment. (b) Distribution of RNA expression and (c) protein marker expression across NSCLC tissues. (d) Visualization of the distribution of AT2 cell subpopulations in adjacent normal and tumor tissues. (e) Comparative UMAP clustering of AT2 cells between adjacent normal and tumor tissuses. (f) Volcano plot illustrating DEGs in AT2 cells between tumor and adjacent normal tissues. (g) Kaplan–Meier survival analysis demonstrating the differences in the survival of patients with high and low expression of the identified DEGs in AT2 cells.

Comparative analyses revealed significant disparities in the composition of AT2 cell subpopulations (**Figure 3d**, **Figure 3e**) between tumor and adjacent normal tissues. Additionally, differentially expressed genes (DEGs) were identified within the AT2 subpopulation of tumor tissues compared with their adjacent normal counterparts (**Figure 3f**). Visualization of the distribution patterns of the four significantly upregulated genes confirmed their predominant expression in tumor tissues (**Figure S6**). The functional significance of these DEGs was explored, revealing pathways involved in the initiation, progression, invasion, metastasis, and therapeutic resistance of lung cancer (**Figure S7**). Patients with high expression levels of these key genes had poor outcomes, indicating their prognostic value (**Figure 3g**). These results increase our understanding of the cellular heterogeneity associated with lung cancer and underscore potential molecular targets and biomarkers for the development of novel therapeutic strategies.

We compared the functional profiles of macrophages and other cell populations in NSCLC by evaluating their activity across key TME-regulating pathways. Significant heterogeneity was observed in signaling pathways activation across various cell types, with tumor-associated pathways being particularly activated in macrophages (**Figure S8**). Further examination revealed marked heterogeneity in the distribution of macrophages between adjacent normal and tumor tissues, emphasizing the variability of distinct cell populations under these two conditions (**Figure S9b**). Comparative analysis of the DEGs in macrophages between adjacent normal and tumor tissues revealed key signaling pathways significantly activated in macrophages within the TME, including those involved in apoptosis, T-cell differentiation, and complement and coagulation cascades, underscoring their potential roles in tumor immune regulation (**Figure S9c-d**). However, functional analysis of these DEGs indicated that their practical utility fell short of expectations (**Figure S10**). Analysis of AT2 cells and macrophages revealed incomplete concordance between the mRNA and protein expression levels, indicating that the transcriptome data are insufficient for assessing cellular states and must be complemented by proteome data. Consequently, TIP-seq, through the integration of proteomic and transcriptomic data, enables more rapid and precise identification of cell populations with high research potential.

### Dendritic Cell (DC) and T-Cell Heterogeneity in NSCLC was Revealed via TIP-seq

Several immune checkpoints that play critical roles in lung cancer progression and the response to immune therapy have been identified. For example, CD276 is associated with immune evasion mechanisms, FGL1 influences the TME and intercellular communication, and IL-4I1 is involved in the regulation of immune responses within tumors. In this work, we employed TIP-seq to analyze both the mRNA and protein expression of these three genes in NSCLC tissues from four new patients (**Table S2**, patients 5-8). By leveraging protein expression data to identify specific cell populations, we investigated the gene expression patterns within these populations to elucidate the roles of specific genes in the pathogenesis of NSCLC.

Nineteen distinct cell populations were identified within the TME (**Figure 4a**). The proportions of these cell populations were compared between adjacent normal and tumor tissues, revealing that epithelial cells, DCs, and T cells were more enriched in lung cancer tissues (**Figure 4b**). Notably, the expression of these three proteins within tumor tissues was predominantly localized to AT2 cells, DCs, and T cells within the TME (**Figure 4d**), indicating considerable discrepancies relative to their corresponding gene expression patterns (**Figure 4c**). The correlation between protein and RNA expression levels was weak, indicating potential posttranscriptional regulation and other factors influencing protein levels (**Figure 4e**). Additionally, this discrepancy was further validated by the complementary expression patterns of these proteins between tumor and adjacent normal tissues (**Figure S11**).

**Figure 4.**
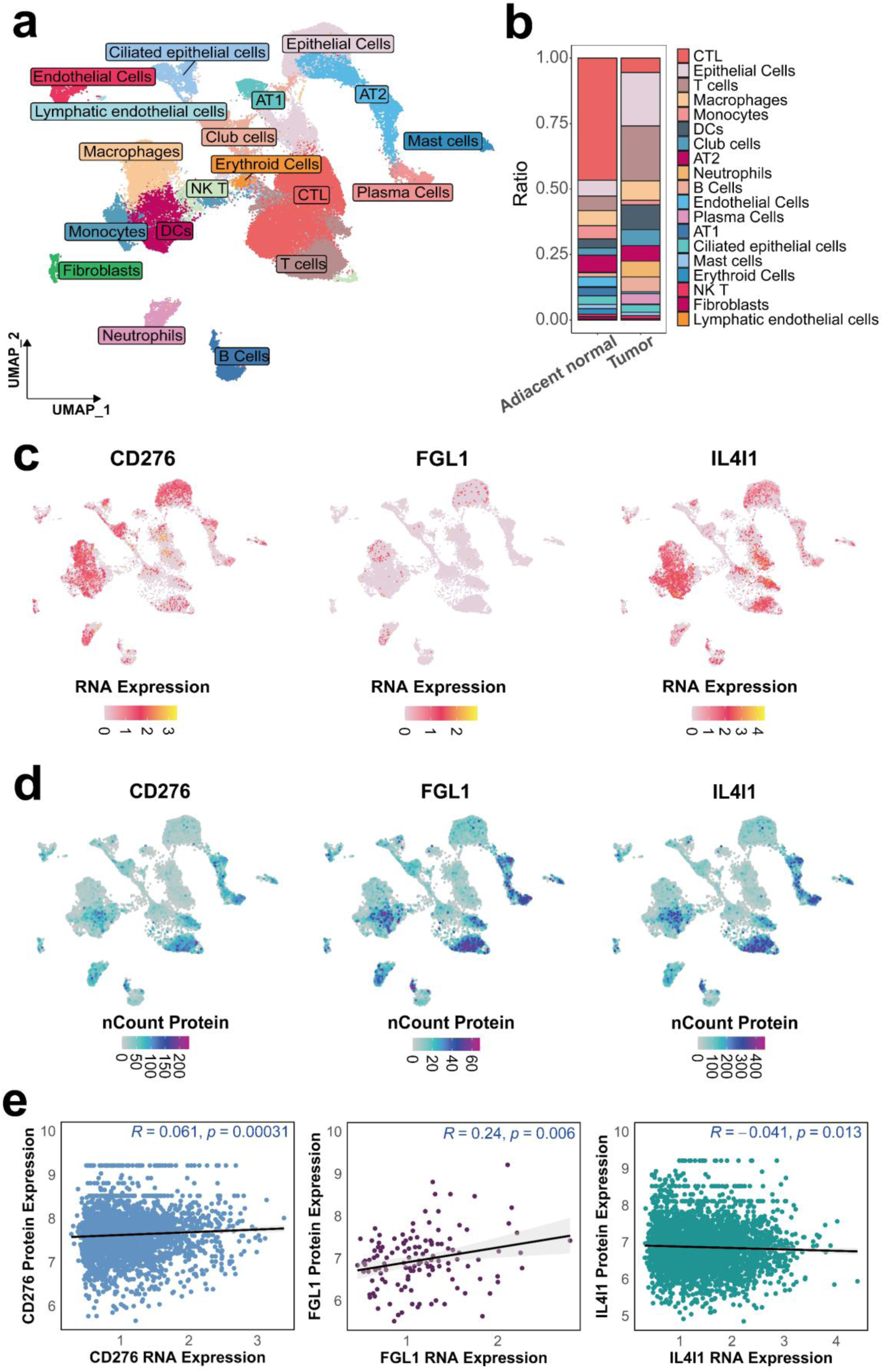
TIP-seq reveals characteristics of NSCLC. (a) UMAP visualization illustrates the cluster assignment. (b) Comparison of the cell populations between adjacent normal and tumor tissues. (c) UMAP plot showing the mRNA and (d) protein expression levels of CD276, FGL1, and IL4I1 across cell subpopulations. (e) Correlation analysis between protein expression and the corresponding RNA expression.

DCs play a vital role in antigen presentation and the response to immune therapy. We investigated the heterogeneity of DC subsets in lung cancer, revealing distinct subpopulation distributions between adjacent normal and tumor tissues, with cDC2s, DC_BAIAP2s, DC_IFI6s and mDCs exclusively detected in tumor tissues, suggesting TME-driven immune cell recruitment (**Figure S12a-b**). Variation in immune cell proportions across patients (**Figure S12c**) revealed interindividual diversity in immune landscapes, implying the potential necessity for patient stratification in DC-targeted therapies. The integrated visualization of gene expression signatures and spatial mapping of DC markers demonstrated functional specialization across different DC subpopulations (**Figure S12d-e**). Differential expression analysis revealed key upregulated and downregulated genes, potentially involved in tumor progression and immune regulation in NSCLC tissues (**Figure S13a**). Immune-related pathways, including those related to complement and coagulation cascades, cytokine‒cytokine receptor interaction, and toll‒like receptor signaling, were significantly enriched in NSCLC tissues (**Figure S13b**). These pathways exhibited distinct activation patterns across DC subpopulations, highlighting their functional specialization in modulating critical immune pathways and suggesting their nonredundant roles in shaping the TME (**Figure S13c-e**). These findings may contribute to the advancement of precision immunotherapies that target specific DC subpopulations.

Building upon these insights, we investigated the heterogeneity of T cells and their potential prognostic implications in lung cancer. The results revealed distinct T-cell clusters in NSCLC tissues and significant compositional differences between tumor and adjacent normal tissues, highlighting their heterogeneity and distribution patterns (**Figure 5a-b, Figure S14a**). T-cell heterogeneity and functional diversity were confirmed, mirroring observations made in DCs (**Figures S14b-c**). Key markers associated with T-cell exhaustion and immune suppression were highly expressed in specific T-cell subpopulations, suggesting functional compromise within the TME (**Figure 5c**). Significant changes in gene expression between tumor and adjacent normal tissues were identified, revealing key DEGs in T-cell populations (**Figure 5d**). Key signaling pathways involved in the immune response and inflammation were enriched (**Figure 5e**). Potential biomarkers related to lung cancer prognosis were identified, with significant differences in the expression of genes such as HSP90AB1, HSPA6, HSPA8 and MUC5B (**Figure 5f**). Patients exhibiting high expression levels of these key genes had poor outcomes, underscoring the prognostic significance of the gene signature (**Figure 5g**). These findings highlight the potential of these genes as therapeutic targets and biomarkers for lung cancer prognosis.

**Figure 5.**
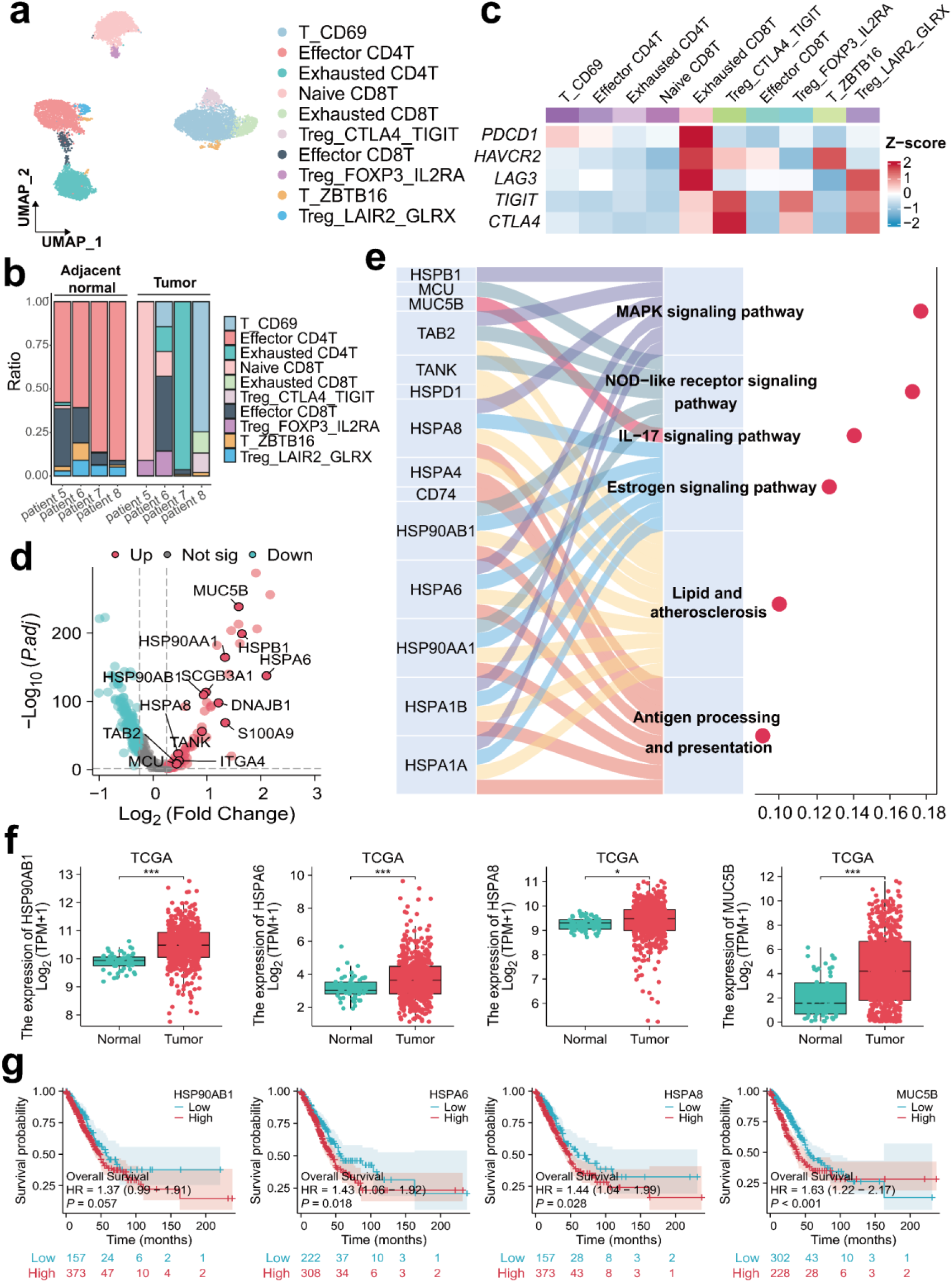
Heterogeneity of T cells and prognostic analysis in lung cancer. (a) UMAP-based visualization of T-cell subpopulations in NSCLC tissues. (b) Proportional analysis of T-cell subpopulations across different samples. (c) Heatmap displaying the expression levels of inhibitory factors across different T-cell subpopulations. (d) Volcano plot showing DEGs between tumor and adjacent normal tissues in T-cell populations. (e) KEGG pathway enrichment analysis of upregulated genes in tumor tissues. (f) Comparison of gene expression levels between adjacent normal and tumor tissues using The Cancer Genome Atlas (TCGA) data. (g) Kaplan‒Meier survival curves showing the impact of high and low expression of key genes on overall survival in lung cancer patients. (* P<0.05, ***P<0.0001)

Furthermore, we analyzed lung squamous cell carcinoma (LUSC) tissue from one patient (**Table S2**, patient 9) and identified significant disparities in protein expression patterns compared with those in LUAD tissue. In LUSC tissue, the three proteins were enriched predominantly in cytotoxic T cells (CTLs) and basal cells (**Figure S15**). There were substantial differences in the basal cell subpopulations between adjacent normal and tumor tissues (**Figure S16a-b**), and the upregulated genes were significantly correlated with poor patient prognosis (**Figure S16c-d, Figure S17**). These findings highlight the distinct molecular characteristics of LUSC and their potential implications for clinical outcomes.

The TIP-seq system liminates the need for individual analysis of each cell subpopulation. By focusing on protein expression, we can efficiently identify relevant cell populations and swiftly identify potential disease-related functional genes from their transcriptomic data. Consequently, the TIP-seq platform holds promise for application in other uncharacterized diseases, facilitating the exploration of innovative strategies for prevention, treatment, and prognosis.

### Functional analysis of tumor-associated immune cells in the TME

To elucidate the interaction between DCs and T cells within the TME of NSCLC, we investigated their functional dynamics and roles in tumor immune regulation. CD8+ T cells in tumor tissues exhibited a relatively high degree of exhaustion, indicating diminished T-cell functionality within the TME (**Figure 6a**). High expression of markers associated with exhausted CD8+ T cells correlated with a poor prognosis in NSCLC patients, underscoring the clinical relevance of T-cell exhaustion in lung cancer (**Figure 6b**). Key ligand‒receptor interactions, such as SPP1‒CD44 and SPP1‒ ITGA4+ITGB1, presented significantly lower p values (p < 0.01) in tumor tissues. These interactions interfere with antigen presentation in immune effector cells and inhibit critical chemokine and cytokine signaling pathways, compromising the efficacy of immune responses (**Figure 6c**). Comparative analysis of immune cell interaction networks revealed increased activity of pathways such as TIGIT in tumor tissues, indicating their potential involvement in immune evasion mechanisms (**Figure 6d**). High expression of MIF, IFN-γ, and TIGIT was correlated with poor prognosis in lung cancer patients, highlighting their potential as therapeutic targets for improving patient prognosis (**Figure 6e**). These findings suggest that combining existing treatments to target these pathways could increase overall survival rates.

**Figure 6.**
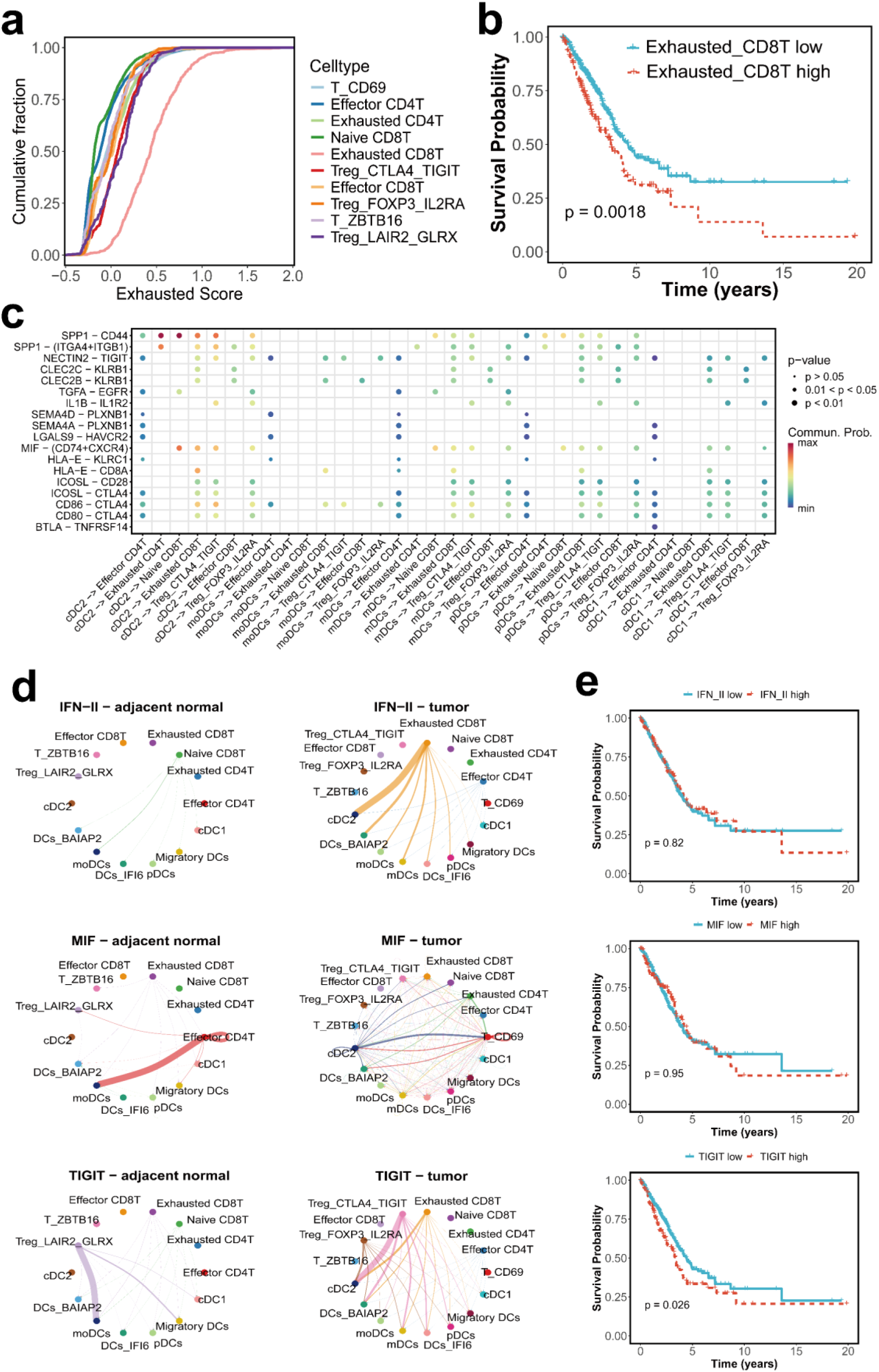
T-cell exhaustion and immune interactions in lung cancer. (a) Exhaustion scores of T-cell subpopulations. (b) Kaplan‒Meier survival analysis showing survival probabilities in NSCLC patients stratified by high and low expression of exhausted CD8+ T-cell markers. (c) Analysis of ligand‒receptor interactions among immune cells. (d) Comparative analysis of immune cell interaction networks. (e) Survival analysis correlating elevated expression of immune pathways members with adverse outcomes in lung cancer patients.

## Discussion

The persistent global burden imposed by lung cancer highlights the urgent necessity for advanced molecular profiling technologies to advance precision cancer treatment^31–33^. While single-cell multiomics has revolutionized our comprehension of tumor heterogeneity^34, 35^, existing methodologies are limited in their ability to analyze transcriptomes and intracellular proteins simultaneously. Currently employed single-cell multiomics platforms face substantial limitations due to two primary constraints: reliance on AOC-based approaches, which can detect only surface or secreted proteins and not intracellular proteins, and perturbations to cellular states resulting from fixation-dependent techniques for intracellular protein detection. In this study, we present TIP-seq, an innovative single-cell multiomics approach that enables the concurrent profiling of transcriptomes and intracellular proteins without necessitating prior cell manipulation.

In contrast to these existing technologies, TIP-seq offers significant advantages in terms of simplicity, cost-effectiveness, and data quality by enabling the simultaneous analysis of transcriptomes and intracellular proteins without prior cell manipulation. By integrating microfluidics and AOC-based approaches, TIP-seq captures mRNA and proteins from individual cells within droplets, thereby achieving high-throughput and precise acquisition of transcriptome and proteome data. In this work, TIP-seq was applied to A549 cell lines and NSCLC tissues, revealing notable cellular heterogeneity and molecular dynamics. Critical immune checkpoint interactions in NSCLC, such as SPP1–CD44, NECTIN2–TIGIT, and CTLA4, alongside functional alterations in tumor-associated DCs and T cells, were identified via TIP-seq. These findings underscore the potential of using this technology to identify novel therapeutic targets and biomarkers in complex diseases. The interaction between T cells and DCs is pivotal for the response to immune therapy, and Itai et al. designed a bispecific dendritic-T-cell engager to facilitate physical interactions between PD-1+ T cells and cDC1s (PMID: 38242085). Bispecific dendritic–T cell engager (BiCE) treatment promoted the formation of active dendritic–T cell crosstalk and resulted in more powerful antitumor immunity than anti-PD1 therapy. Through TIP-seq, we revealed that exhausted CD8+ T cells, exhausted CD4+ T cells, and Tregs interact with DCs in lung cancer tissues, which may provide valuable information for the development of novel BiCEs. While TIP-seq presents notable advantages, it is not without its limitations. Like other methodologies, it depends on antibodies for protein capture, which inherently constrains its accuracy and sensitivity on the basis of the affinity and availability of these antibodies. Additionally, the speed and efficiency of droplet encapsulation may lead to incomplete or uneven protein capture. Moreover, as the volume of single-cell data continues to expand, the complexity of data analysis correspondingly increases, and existing computational tools may no longer be adequate for the analysis of these generated data.

Moving forward, TIP-seq has demonstrated considerable potential in cancer research and beyond. Furthermore, the integration of TIP-seq with other single-cell technologies, such as spatial transcriptomics and single-cell metabolomics, could facilitate multidimensional single-cell analysis, offering a more comprehensive perspective in cancer research. In conclusion, TIP-seq holds considerable promise for advancing precision medicine and clinical applications within cancer research, while simultaneously opening new pathways for the advancement of cell-based therapies and regenerative medicine.

## Materials and Methods

### Chemicals and Reagents

All oligonucleotides were synthesized by Sangon Biotech (Shanghai, China). Capture and detection antibodies against CEA, Cyfra21-1, NSE, proGRP, and SCCA were purchased from AOKEIBOTAI (Wuhan, China), whereas antibodies targeting CD276, FGL-1, and IL-4I1 were sourced from Abcam (UK).

### Antibody-Oligonucleotide Conjugates

Initially, 100 µg of detection antibody was diluted to a concentration of 2 mg/mL in PBS and incubated with 2 µL of 10 mM NHS–PEG4–N3 (Thermo Scientific, 26130) at room temperature for 2 hours to facilitate the introduction of azide groups onto the antibody (**Figure S1a**). Following this reaction, unreacted NHS–PEG4–N3 was removed by performing three successive washes with Amicon Ultra-0.5 mL 50 kDa centrifugal filters. The antibodies were then transferred to a tube, and 20 µL of 100 µM DBCO-modified oligonucleotide was added. The total volume was adjusted to 100 µL with PBS, and the mixture was incubated at 4℃ for 16 hours to achieve covalent conjugation. An equal volume of saturated ammonium sulfate solution was added to the reaction mixture, followed by incubation on ice for 20 minutes. The mixture was centrifuged at 15,000 relative centrifugal force (rcf) for 5 minutes, after which the supernatant was discarded, and the pellet was redissolved in PBS to obtain the AOC^36, 37^. As shown in **Figure S1b** and **Figure S1c**, the leftward shift of the absorption peak and the upward shift of the bands in the gel image both indicate successful conjugation of the oligo to the antibody.

### Antibody-coated Beads Preparation

The conjugation process of the capture antibodies with NHS-Biotin (Thermo Scientific, A35358) parallels that of the detection antibodies with NHS–PEG4–N3. A total of 100 µg of antibody‒biotin conjugates was mixed with 100 µL of streptavidin-coated Dynabeads (Thermo Scientific, 65305) and incubated at ambient temperature for 2 hours. Subsequent magnetic separation facilitated the removal of unbound antibodies, yielding antibody-coated beads.

### Preparation of Barcoding Hydrogel Beads

Eight milligrams of BAC was fully solubilized in 300 µL of DMSO through vigorous vortexing, followed by the addition of 700 µL of H_2_O and thorough mixing. The resulting mixture was then incubated at 60℃ in a shaker for 2 hours to yield a 0.8% BAC solution. For the preparation of droplets using a microfluidic chip, the final solution of 1 mL consisted of the following: 100 µL of TBSET buffer, 60 µL of 10% APS, 150 µL of 40% acrylamide solution, 100 µL of 100 µM acrylamide-modified oligo, 490 µL of 0.8% BAC solution, and 100 µL of H_2_O. The droplets were incubated at 65℃ for 14 hours to facilitate polymerization into hydrogel beads. Demulsification was subsequently carried out using 20% perfluorooctanol, and any residual oil phase was eliminated by washing with 1% Span 80 in hexane. The hydrogel beads underwent three washes in the washing buffer, with each wash followed by centrifugation at 3,000 rcf for 3 minutes to collect the beads.

For the ligation reaction, a ligation premix solution, consisting of 8 mL of hydrogel beads, 800 μL of 100 μM complementary oligonucleotide, 16,960 μL of H_2_O, 2,320 μL of 10× ligation buffer, and 1,520 μL of 400 U/μL T4 ligase was prepared. Subsequently, 37 μL of the premix was aliquoted into each well of a plate, to which 2 µL of 66 µM barcode1 was added. The plate was vortexed to ensure uniformity, sealed, and incubated overnight at room temperature on a rotating mixer. To terminate the ligation reaction, 100 µL of stop buffer (50 mM Tris-HCl, 50 mM EDTA, 0.1% Tween 20) was added to each well. The hydrogel beads were then recovered by centrifugation at 3,000 rcf for 3 minutes, and the supernatant was discarded. The beads were subjected to an additional three washes with washing buffer. Barcode2 ligation was performed using an identical protocol.

### A549 Cell Culture

The A549 cell line, sourced from the Chinese Typical Culture Collection (CTCC), was cultured in DMEM supplemented with 10% (v/v) fetal bovine serum (FBS) and 1% penicillin/streptomycin (P/S). The cells were incubated at 37°C in a 5% CO_2_ environment.

### Enzyme-Linked Immunosorbent Assay

Initially, the capture antibody was diluted to a concentration of 5 μg/mL and applied to the ELISA plate overnight at 4℃. The plate was subsequently washed three times with 0.05% PBST. The standards and samples were then introduced and incubated for 2 hours at room temperature. Following another set of three washes, the plate was blocked with 0.5% BSA in PBS for 2 hours. The biotinylated detection antibody was then added, and the plate was incubated for 1 hour at room temperature, followed by six washes. Next, the plate was incubated with streptavidin-horseradish peroxidase (SA-HRP) (Thermo Scientific, S911) for 30 minutes, washed eight times, and treated with tetramethylbenzidine (TMB) (YEASEN, China, 36602ES60) for 15 minutes. The reaction was terminated by the addition of an ELISA stop solution, and the optical density (OD) was measured at 450 nm. Sample concentrations were determined by interpolation from the standard curve.

### Preparation of Single-Cell Suspension from Lung Cancer Tissues

The tissue was thoroughly rinsed with PBS and transferred to a culture dish, where it was thoroughly minced into the smallest possible fragments. The tissue fragments were subsequently subjected to enzymatic digestion using a tumor tissues dissociation kit (RWD, DHTEH-10) at 37℃ for 30 minutes. After digestion, the resulting single-cell suspension was filtered through a 30 μm cell strainer into a 15 mL centrifuge tube and maintained on ice. For any remaining undigested tissue, 1 mL of trypsin was added, and the digestion process was extended for an additional 15–20 minutes at 37℃. The dissociated single cells were then filtered and pooled with the initially isolated cells. The combined cell suspension was centrifuged at 300 rcf for 5 minutes to pellet the cells. Next, 2 mL of red blood cell lysis buffer was added to the cell pellet, and the mixture was gently pipetted to ensure thorough resuspension. The suspension was incubated at room temperature for 5–10 minutes, after which 12 mL of PBS was added to quench the lysis reaction. The cells were pelleted again by centrifugation at 300 rcf for 5 minutes and finally resuspended in PBS for further use.

### Microfluidic Device Fabrication

The microfluidic chip was designed using AutoCAD 2021(Autodesk, USA) to enable hydrogel bead generation (**Figure S2a**) and cell encapsulation (**Figure S2c**). The device molds were fabricated using standard photolithography, where SU-8 photoresist (Kayaku Advanced Materials, Japan, Kayaku SU-8 2050) was spin-coated onto silicon wafers and patterned by UV lithography using an MA/BA6 Gen4 aligner (SUSS MicroTec, Germany). To ensure the easy release of polydimethylsiloxane (PDMS), the SU-8 mold was silanized by overnight exposure to trichloro (1H,1H,2H,2H-perfluorooctyl) silane (Sigma Aldrich, USA, 448931) vapor under vacuum.

A 10:1 (w/w) mixture of PDMS prepolymer and curing agent (Sylgard 184 Dow Corning, USA, 01673921) was prepared, degassed for 30 min, poured onto the mold, and cured at 80°C for 30 min. After demolding, inlet and outlet ports were created using a biopsy punch (Electron Microscopy Sciences, USA, 2023020303). The PDMS replica was bonded to a glass slide via oxygen plasma treatment (Electronic Diener, Atto, Germany, 18 W, 30 s). Finally, the microfluidic channels were rendered hydrophobic by trichlorosilane treatment to prevent aqueous adhesion. The assembled chips were stored in a desiccator until use.

### Cell Encapsulation

First, the cell encapsulation microfluidic chip was positioned on the stage of an inverted microscope (Nikon Ti2). A tygon hose was subsequently connected to the chip’s channel and linked to an air pump. Next, the cell mixture, gel beads, and oil were introduced into the microfluidic device. The air pump was then activated, with the cell mixture and barcoded bead mixture each maintained at a pressure of 70 mbar, while the oil pressure was set to 150 mbar. Upon completion of the cell encapsulation process, the emulsified droplets at the outlet were collected into a tube and sealed with mineral oil.

### Single-cell Dual-omics Library Construction and Data Analysis

Following the coencapsulation of cells and hydrogel beads and subsequent reverse transcription, the droplets underwent demulsification with 20% perfluorooctanoic to eliminate the oil phase. The supernatants, which contained transcriptomic data, and the beads, which contained proteomic information, were then separated through magnetic isolation. Library preparation for both the transcriptome and proteome was performed using a LeadOmics single-cell 3’ RNA kit (Lead Healthcare, China, 10401-001419).

Differentially expressed gene (DEG) analysis for each cell cluster was performed using the FindMarkers function in Seurat (v4.1.0), employing the Wilcoxon rank-sum test (PMID:36973787). DEGs were filtered on the basis of the following criteria: expression in more than 25% of the cells within the cluster, a log2-fold change (FC) exceeding 0.25, and a false discovery rate (FDR) below 0.05.

The Wilcoxon rank-sum tests were conducted using the FindMarkers function to identify significant tumor-related genes. Volcano plots were constructed to visualize DEGs whose log2 FC was greater than 1 or less than -1 and whose q value was less than 0.05. Functional enrichment analysis was performed via clusterProfiler (v4.2.2, PMID:2245546), and the expression patterns of DEGs across cell subpopulations were examined.

CellChat (v1.1.3, PMID:35938159) was utilized to investigate cell‒cell communication within the TME. The data preprocessing steps involved several steps, including the identification of overexpressed genes and interactions, as well as data projection, to assess the strength of ligand‒receptor (L‒R) interactions and the communication probabilities among cell clusters. Further analysis of receptor‒ligand interactions was conducted using CellPhoneDB (v2.1.4, PMID:32103204).

Pseudotime trajectory analysis was performed using Monocle (v2.22.0, PMID:28825705). The normalized and annotated single-cell expression matrix served as the input, with highly variable genes being selected and genes with low expression being excluded. Dimensionality reduction was achieved using the ReduceDimension function with the DDRTree method to construct pseudotime trajectories.

## Data Availability

The data that support the findings of this study have been deposited into the CNGB Sequence Archive (CNSA) of the China National Genebank Database (CNGBdb) with accession number CNP0007354.

## Acknowledgments

We acknowledge the financial support from the Major Project of Guangzhou National Laboratory (grant no. GZNL2024A03001) and the National Natural Science Foundation of China (grant no. 82202040).

---

**Figure S1.**
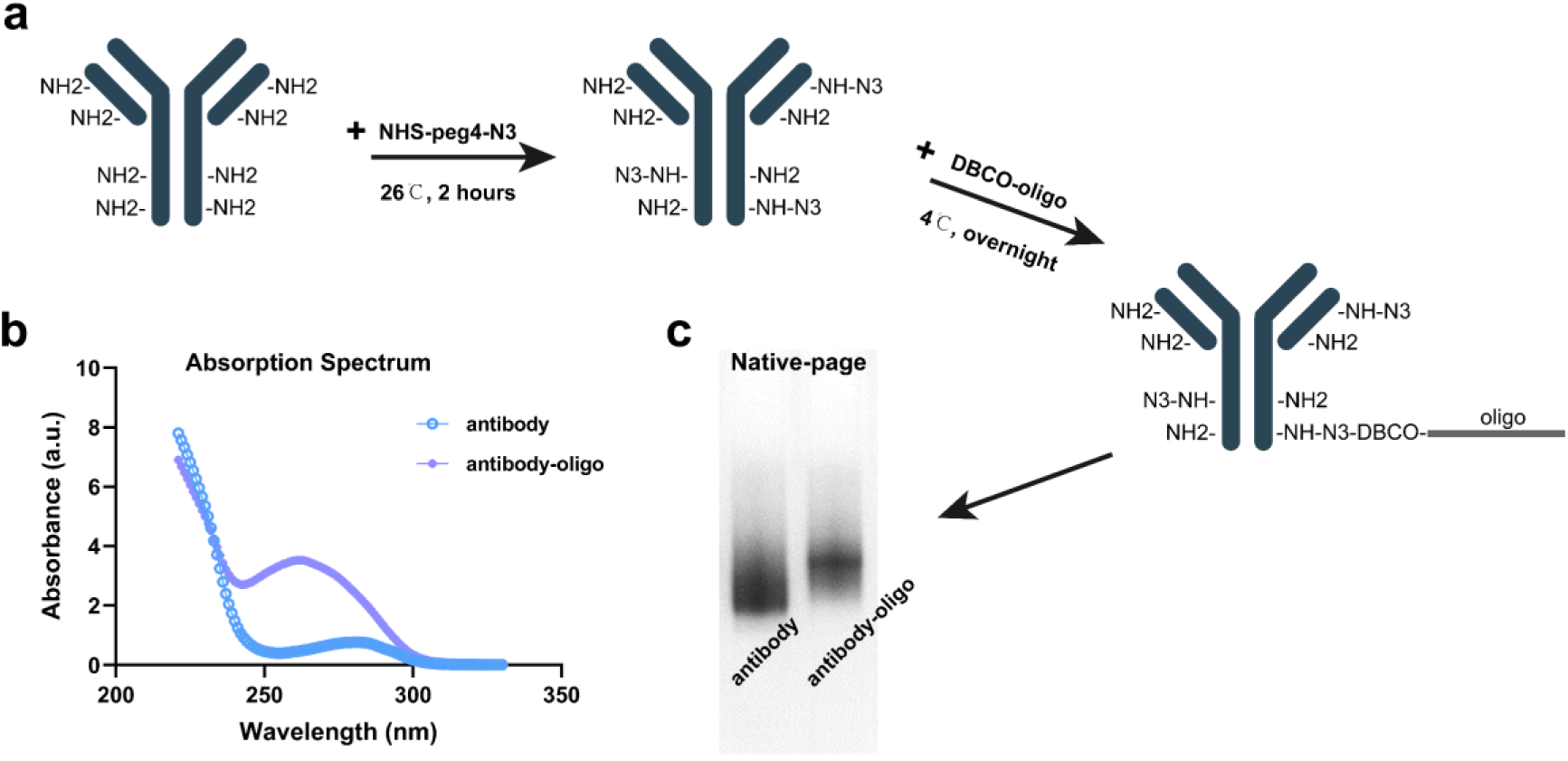
(a) A schematic representation illustrating the conjugation process of DBCO-modified oligonucleotides and antibodies. (b) The observed shift in the absorption peak toward 260 nm serves as evidence for the successful conjugation of oligonucleotides to the antibodies. (c) A comparative native gel image depicting the AOCs alongside unmodified antibodies.

**Figure S2.**
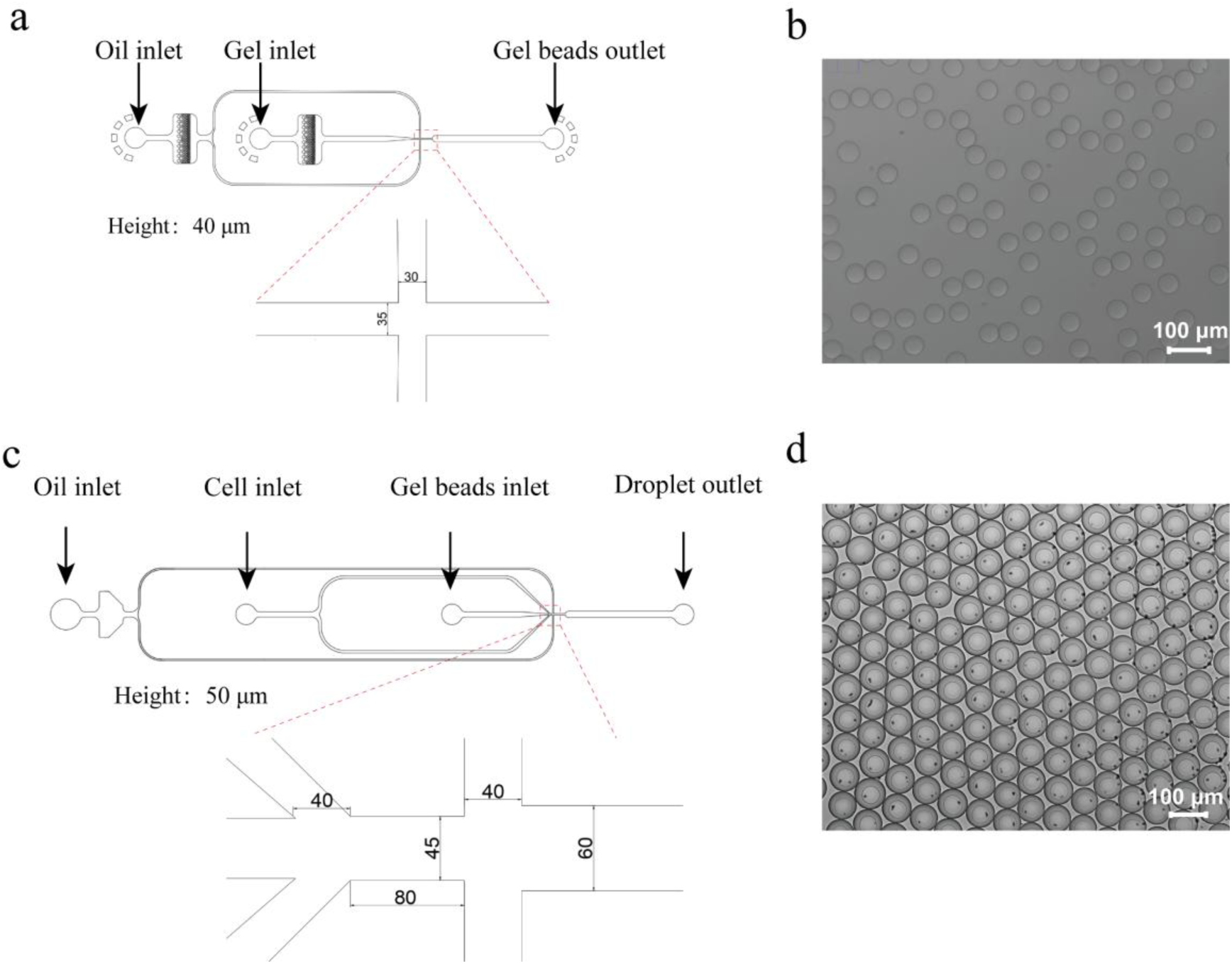
(a) Schematic representation of the microfluidic chip designed for hydrogel bead generation. Height: 40 μm. (b) Microscopy image of the hydrogel beads. Scale bar:100 μm. (c) Schematic representation of the microfluidic chip designed for cell encapsulation. Height: 50 μm. (d) Microscopy image of the encapsulated droplets. Scale bar:100 μm.

**Figure S3:**
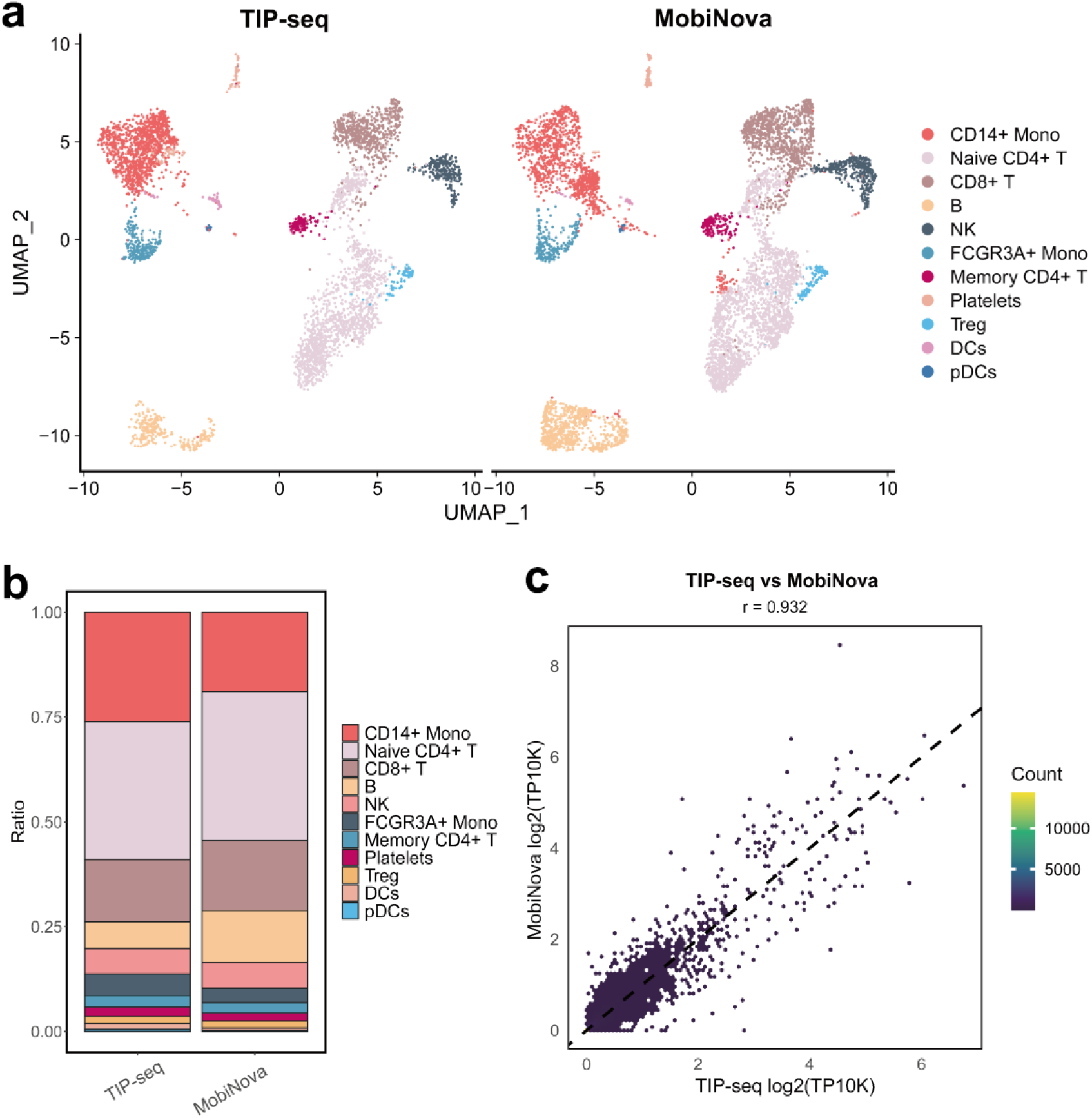
Comparative analysis of TIP-seq and MobiNova in PBMC clustering. (a) Comparative evaluation of cell clustering outcomes between TIP-seq and 10x Genomics in PBMC samples. (b) Assessment of the ability of the two technologies to detect cell proportions. (c) Correlation analysis between the two methodologies.

**Figure S4.**
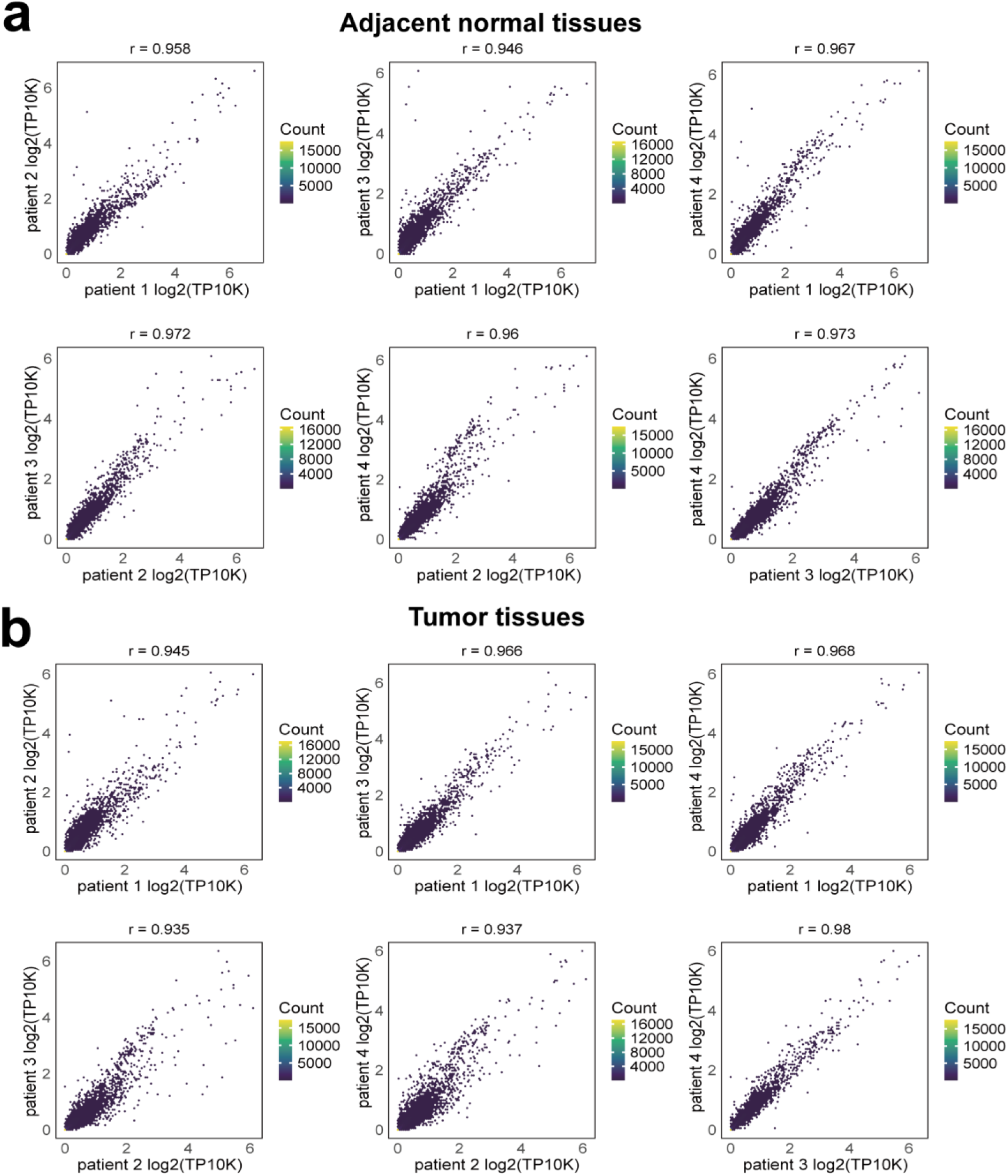
Analysis of transcriptomic correlation across NSCLC samples. (a) Correlation analysis of adjacent normal tissues from patients 1-4. (b) Correlation analysis of tumor tissues from patients 1-4.

**Figure S5.**
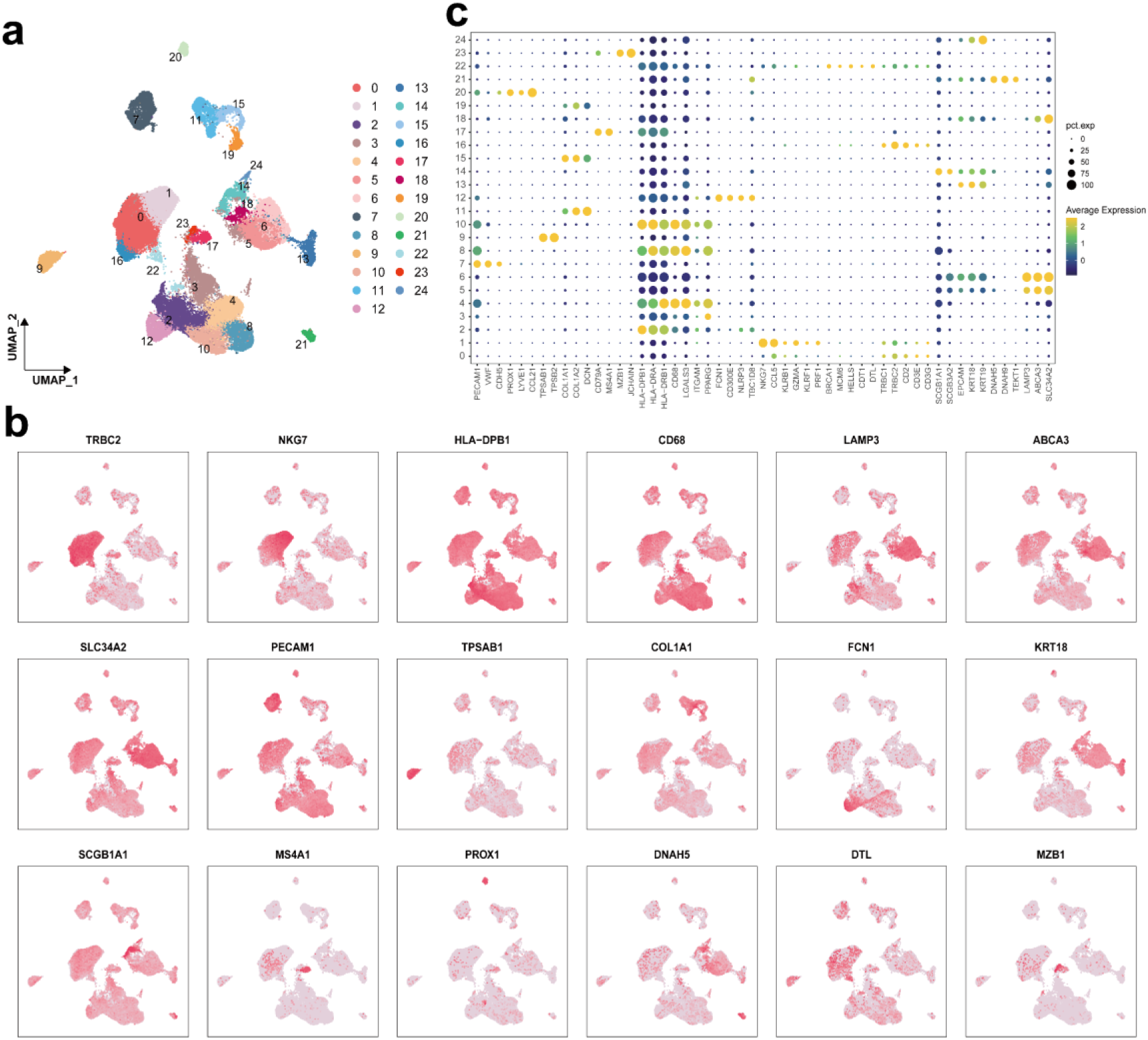
Single-cell transcriptomic profiling of NSCLC tissues. (a) UMAP visualization of cell clusters identified in NSCLC tissues. (b) Dot plot displaying the expression levels of key genes across various cell clusters, where the size of each dot represents the percentage of cells expressing the gene and the color intensity reflects the expression level. (c) Expression patterns of marker genes in their respective cell clusters, demonstrating cell-type-specific gene signatures.

**Figure S6.**
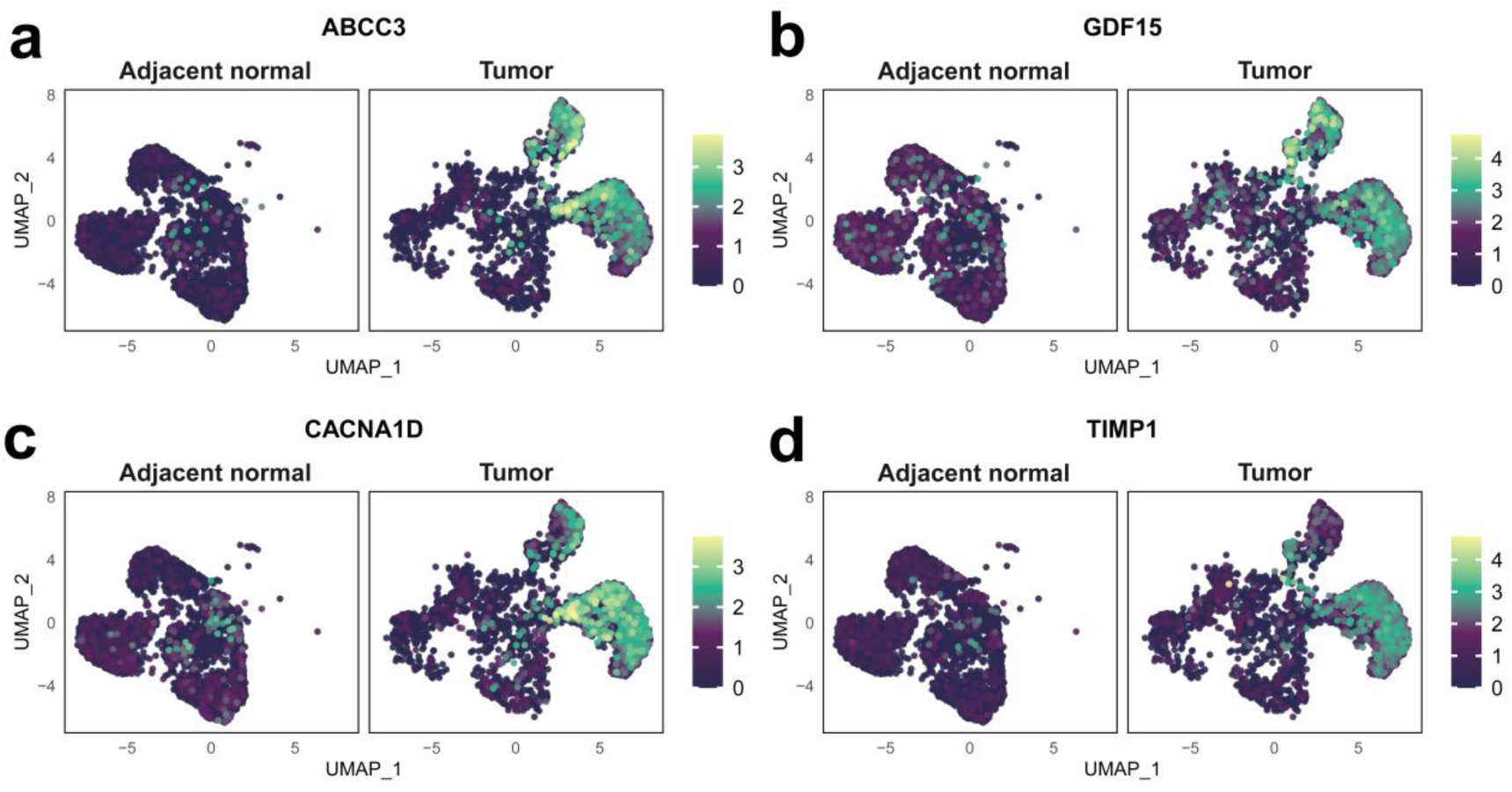
Comparative distribution of the four genes whose expression was upregulated in AT2 cells between adjacent normal and tumor tissues. (a) ABCC3: ATP binding cassette subfamily C member 3. (b) GDF15: growth differentiation factor 15. (c) CACNA1D: calcium voltage-gated channel subunit alpha1 D. (d) TIMP1: tissue inhibitor of metal protease 1.

**Figure S7.**
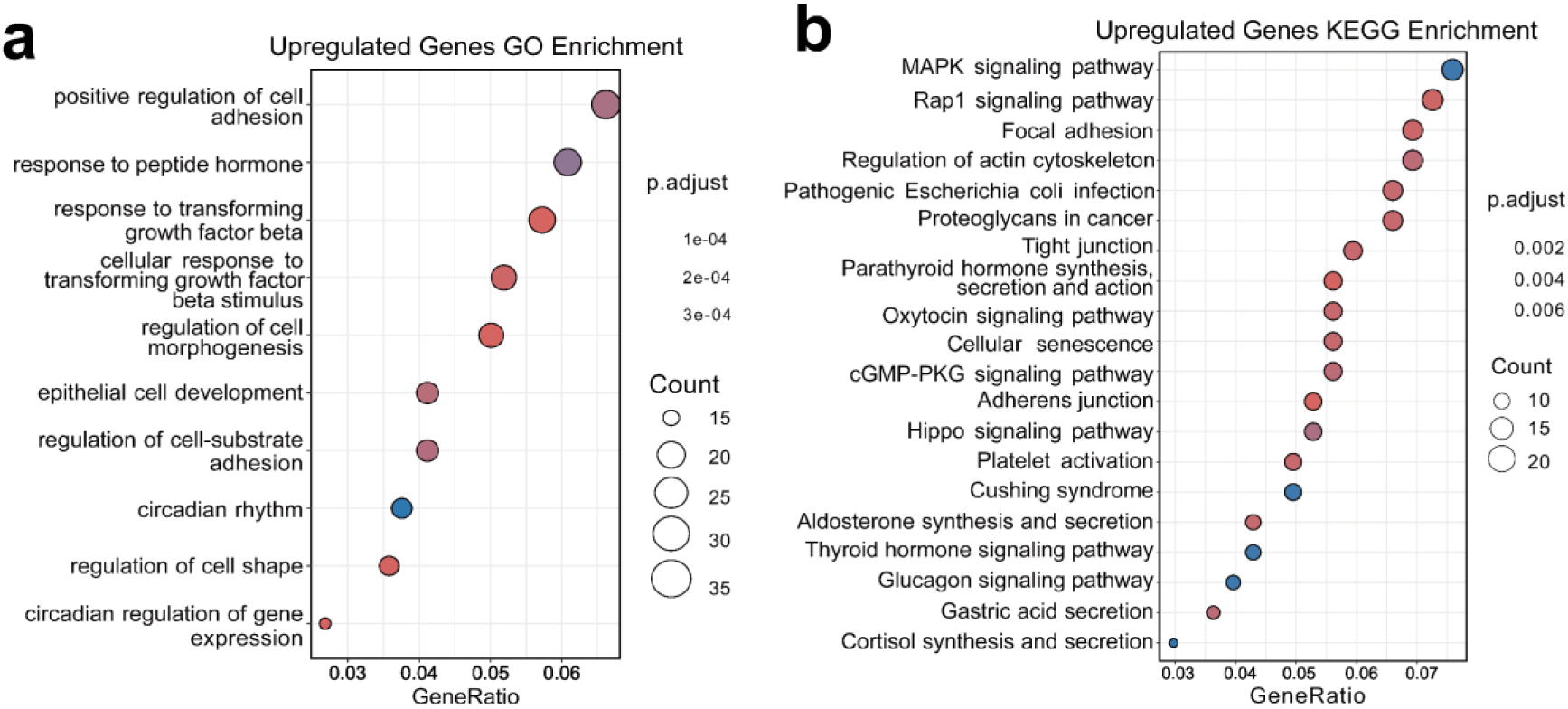
(a) GO functional enrichment analysis of biological processes associated with the upregulated genes in AT2 cells. (b) KEGG pathway enrichment analysis of the upregulated genes in AT2 cells.

**Figure S8.**
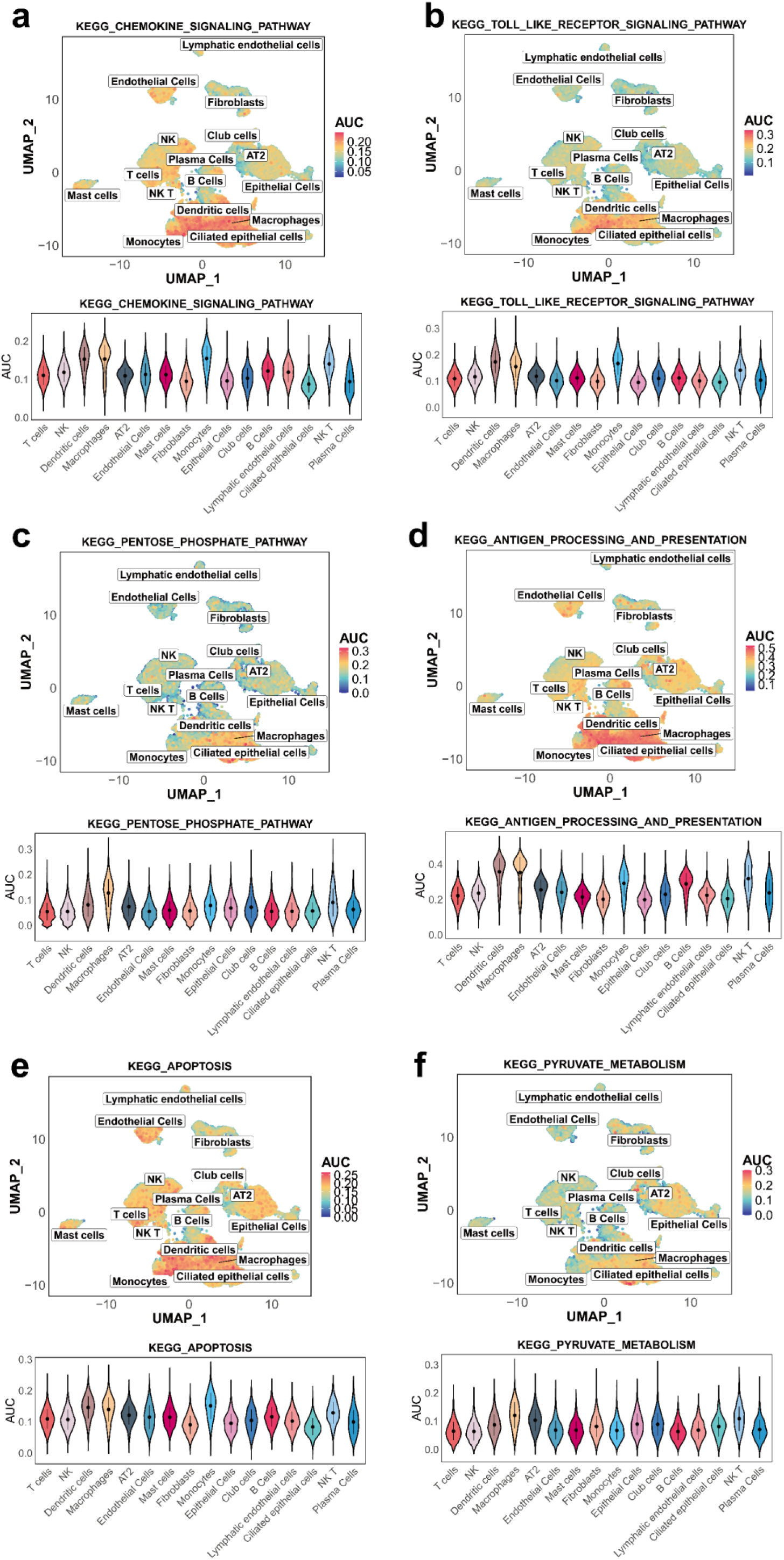
UMAP and violin plots were used to evaluate the expression patterns and diagnostic utility of key signaling pathways across various cell types in tumor and adjacent normal tissues. The violin plots illustrate the area under the curve (AUC) values, which indicate the diagnostic potential of these pathways, with a particular emphasis on macrophages. The pathways analyzed included: (a) chemokine signaling-related pathways, (b) the toll-like receptor signaling pathway, (c) the pentose phosphate pathway, (d) pathways associated with antigen processing and presentation, (e) pathway related to apoptosis, and (f) pyruvate metabolic pathways.

**Figure S9.**
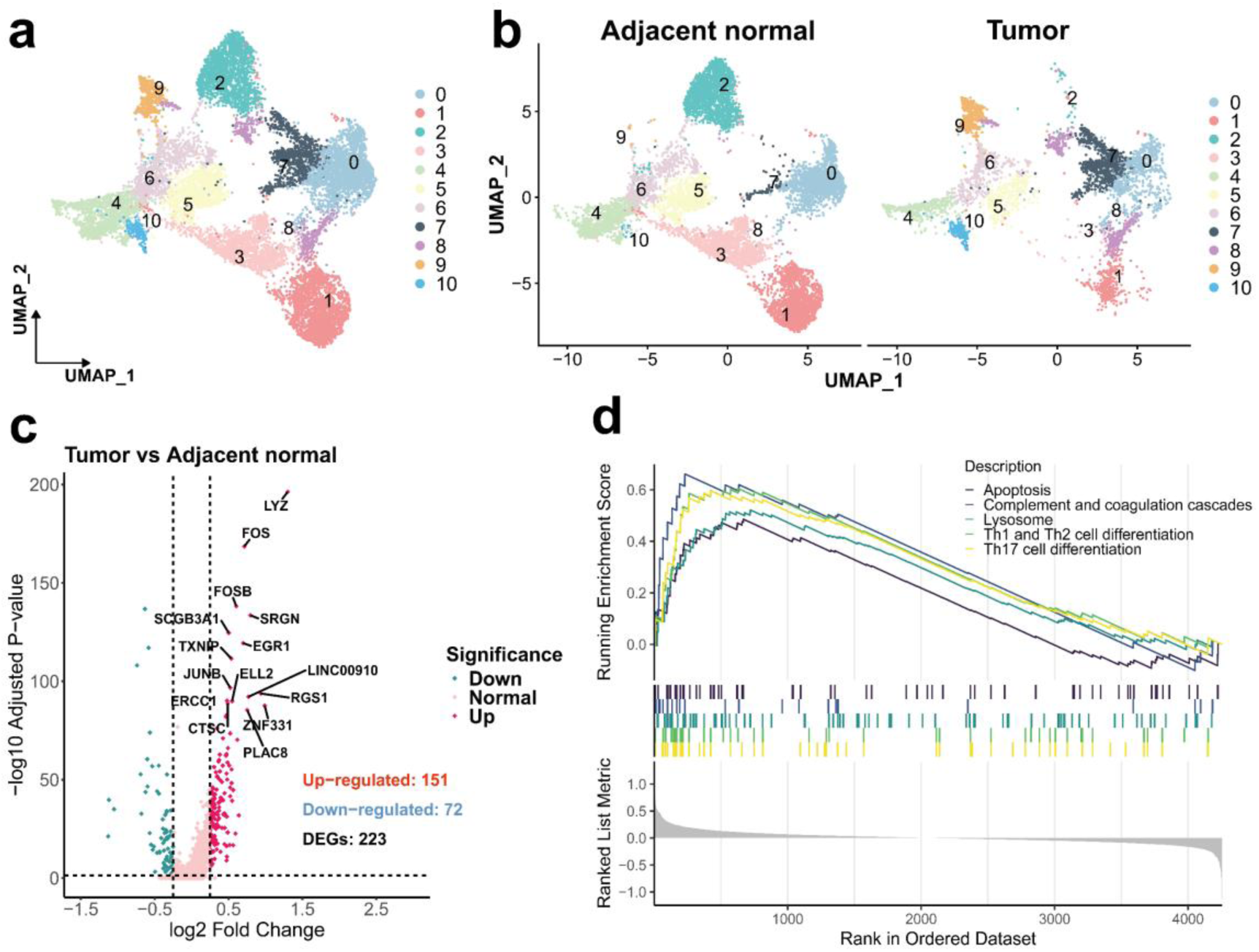
Analysis of macrophage traits in LUAD samples. (a) UMAP visualization showing macrophage clusters. (b) Comparative analysis of macrophage distribution in adjacent normal versus tumor tissues. (c) Analysis of gene expression differences in macrophages. (d) Gene set enrichment analysis (GSEA) of the principal signaling pathways activated in macrophages within the TME.

**Figure S10.**
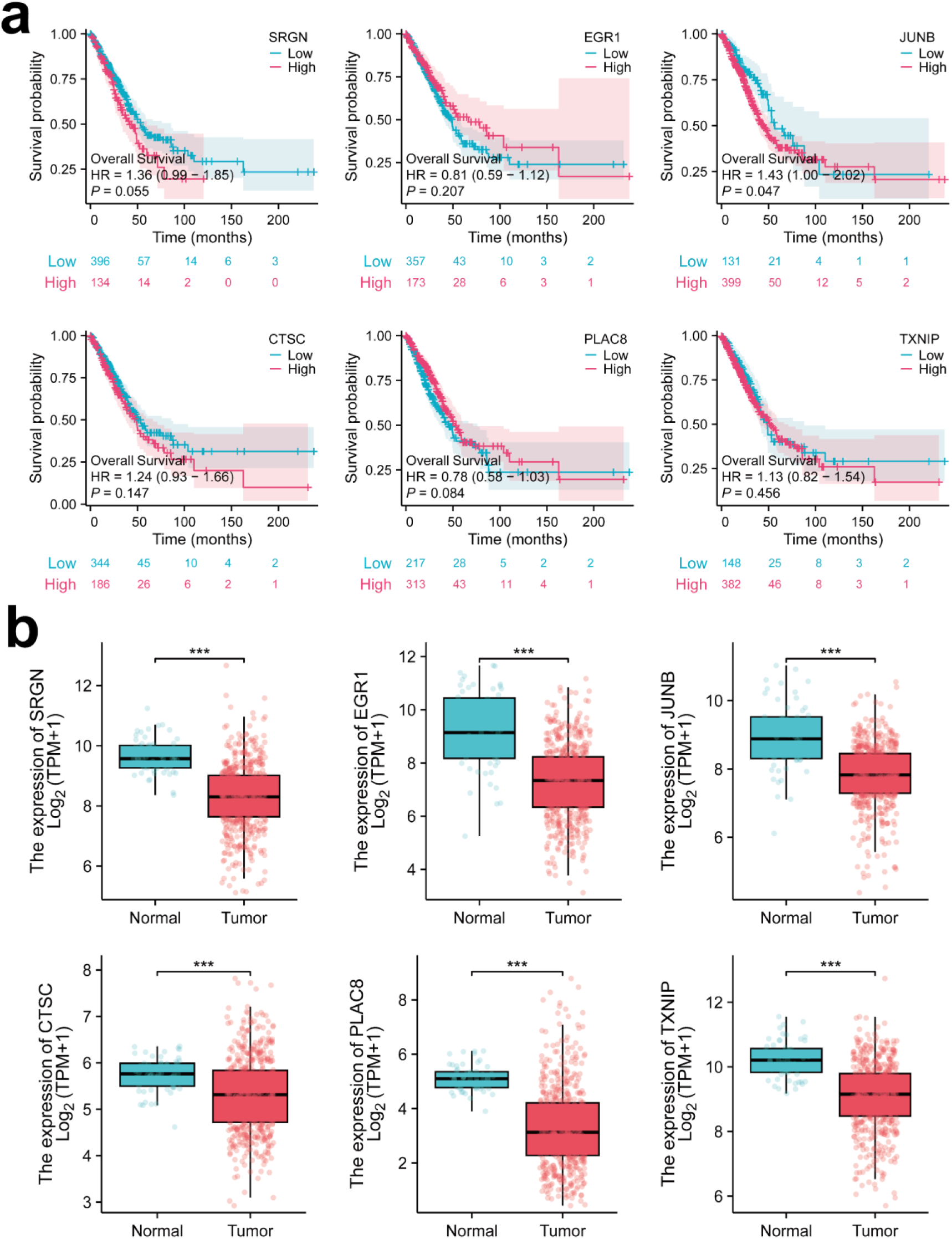
(a) Kaplan‒Meier survival analysis showing the survival disparities between cohorts with high and low expression of the identified differentially upregulated genes in macrophages. (b) Comparative analysis of genes with elevated expression levels in macrophages on the basis of data derived from the TCGA.

**Figure S11:**
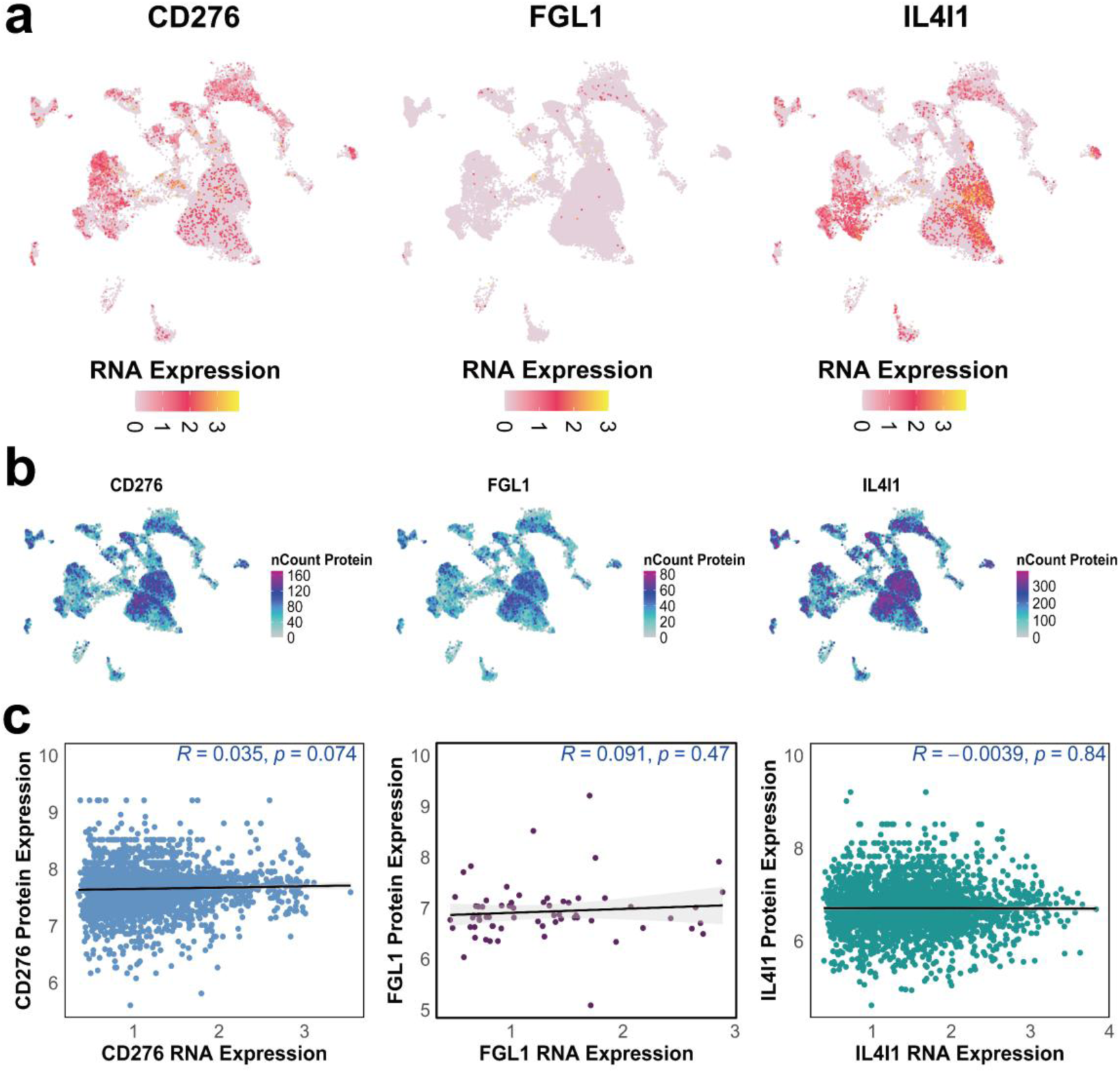
(a) Distribution of the RNA expression levels of CD276, FGL1, and IL4I1 in adjacent normal tissues. (b) Corresponding distribution of protein expression levels. (c) Scatter plots depicting the correlation between RNA and protein expression levels for each gene.

**Figure S12.**
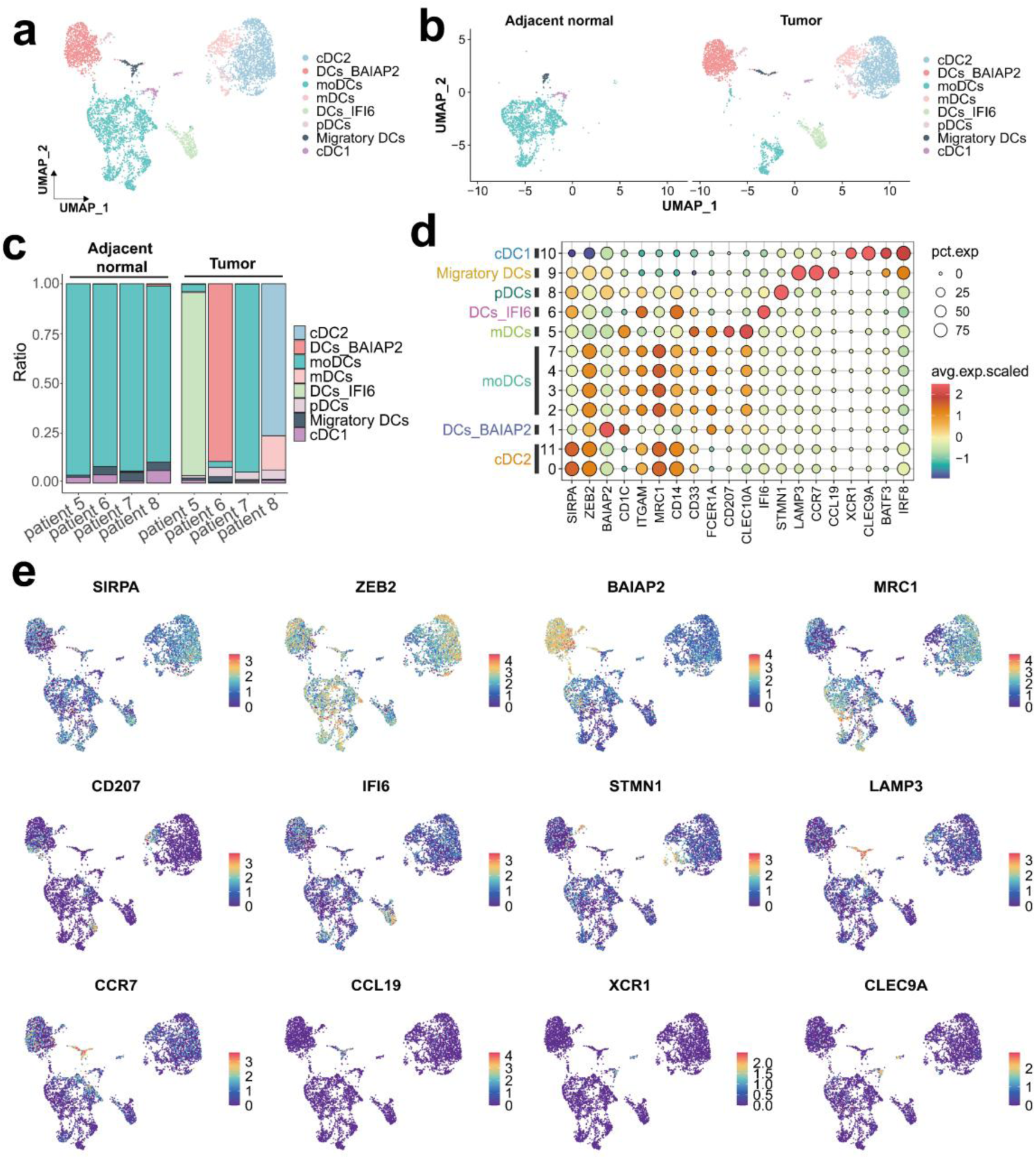
Heterogeneity of dendritic cell subpopulations in lung cancer. (a) UMAP visualization of DC subpopulations in lung cancer tissues, revealing distinct clusters and heterogeneity. (b) Comparison of DC subpopulation distributions between normal and lung cancer tissues, showing enrichment of certain subsets in tumors. (c) Proportional analysis of DC subpopulations across different samples, highlighting interindividual heterogeneity in the immune microenvironment. (d) Dot plot displaying distinct expression profiles of DC subpopulation-specific genes, indicating functional diversity. (e) UMAP visualization of the spatial distribution of marker genes in DC subpopulations, highlighting functional diversity within the TME.

**Figure S13.**
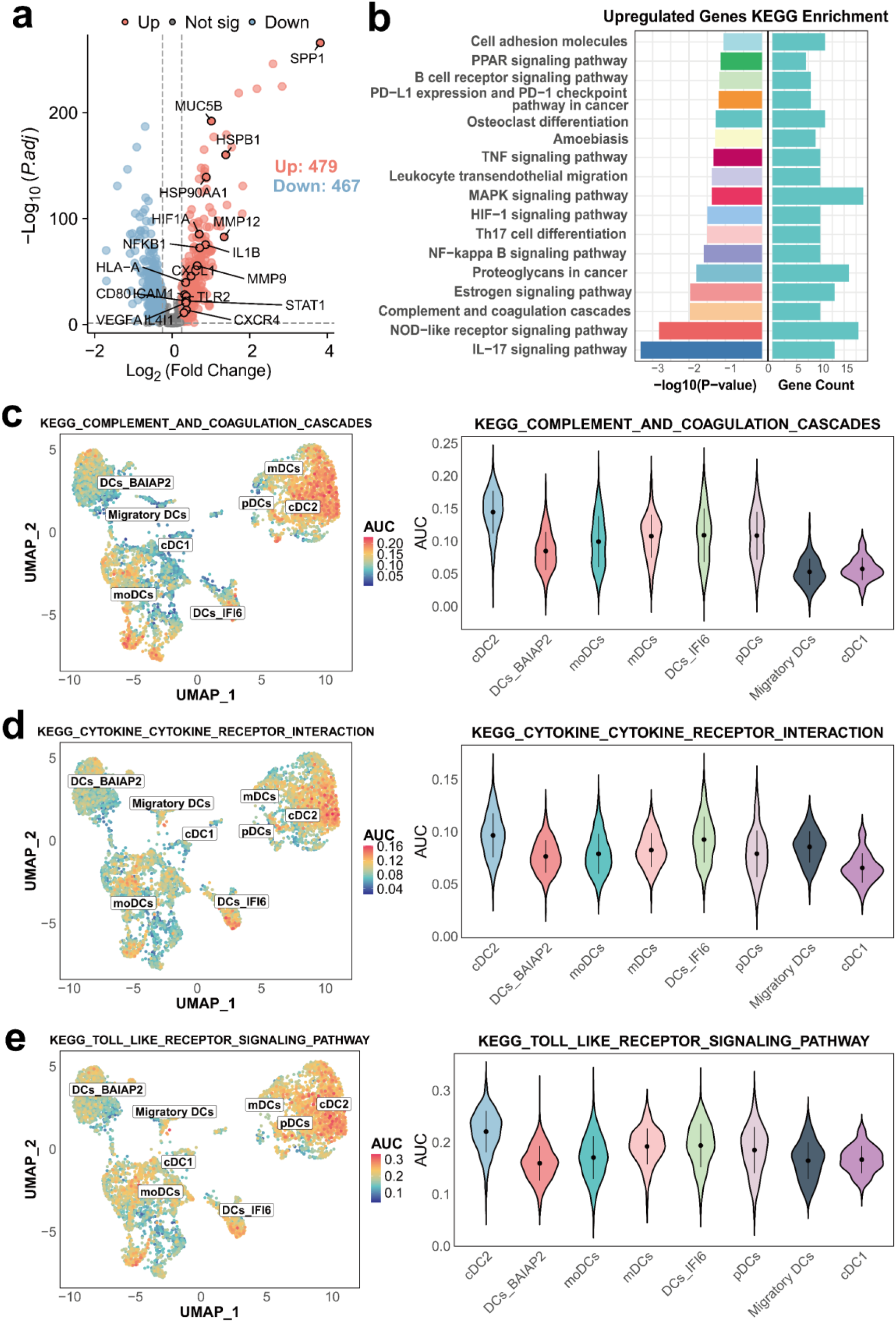
Differential expression analysis reveals immune-related signaling pathway activation in lung cancer. (a) Volcano plot showing genes that are differentially expressed between lung cancer and normal tissues, highlighting key upregulated and downregulated genes. (b) KEGG analysis revealed significant enrichment of immune-related pathways, including pathways associated with complement and coagulation cascades, cytokine‒cytokine receptor interaction, and toll‒like receptor signaling, in lung cancer tissues. (c) UMAP visualization of complement and coagulation cascade pathway activation in different DC subpopulations, with AUC scores used to quantify pathway activity levels. (d) UMAP visualization of cytokine‒cytokine receptor interaction pathway activation in different DC subpopulations, highlighting distinct activation patterns. (e) UMAP visualization of toll‒like receptor signaling pathway activation in different DC subpopulations, with AUC scores indicating pathway activity levels.

**Figure S14:**
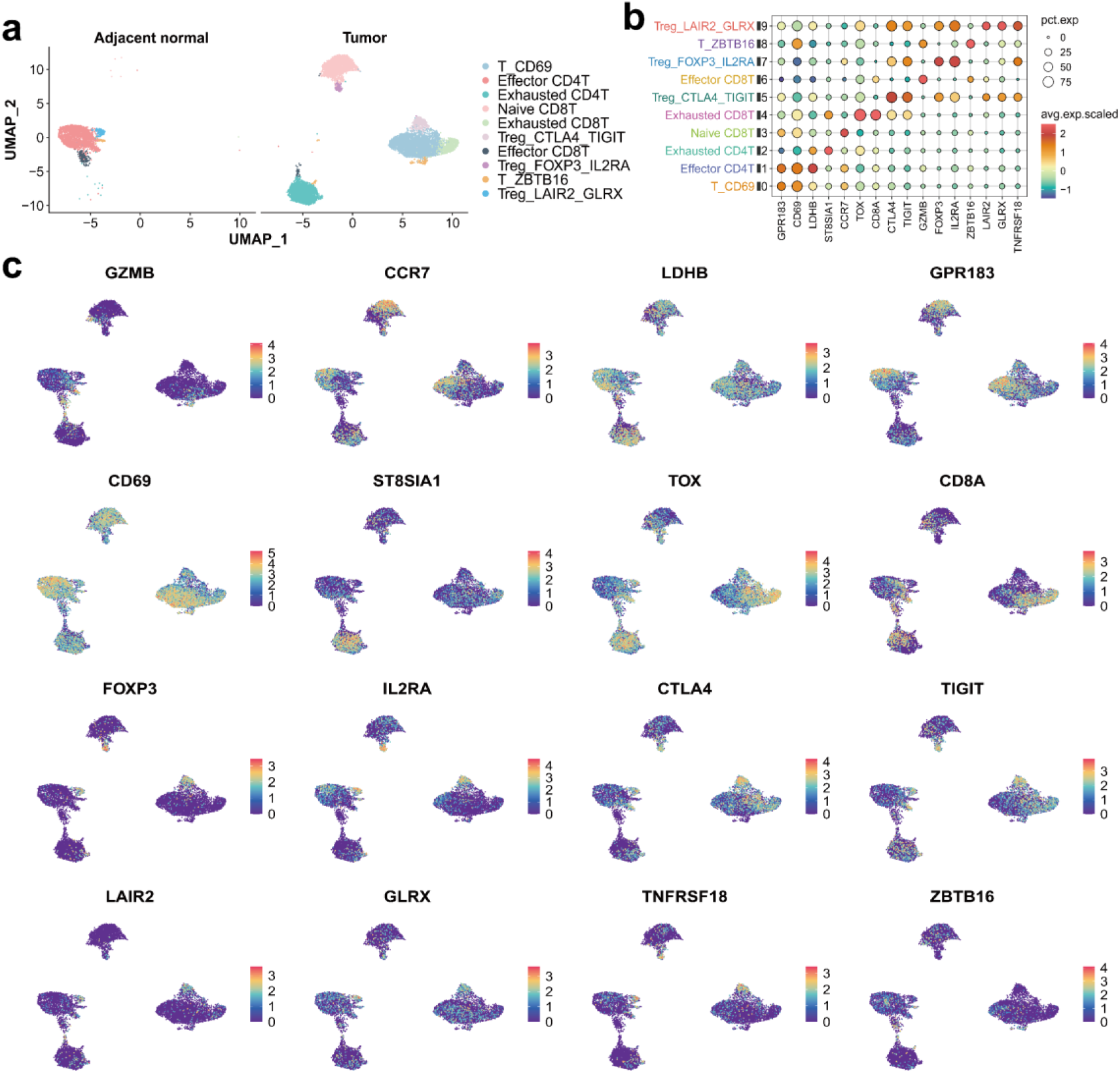
T-cell subpopulation heterogeneity and key gene expression in lung cancer. (a) UMAP visualization of T-cell subpopulations in normal and lung cancer tissues, revealing distinct clusters and heterogeneity. (b) Dot plot showing the expression abundance and proportion of functional genes in T-cell subpopulations, highlighting distinct profiles and functional diversity. (c) UMAP visualization of spatial expression patterns of marker genes in T-cell subpopulations, highlighting functional diversity within the TME.

**Figure S15.**
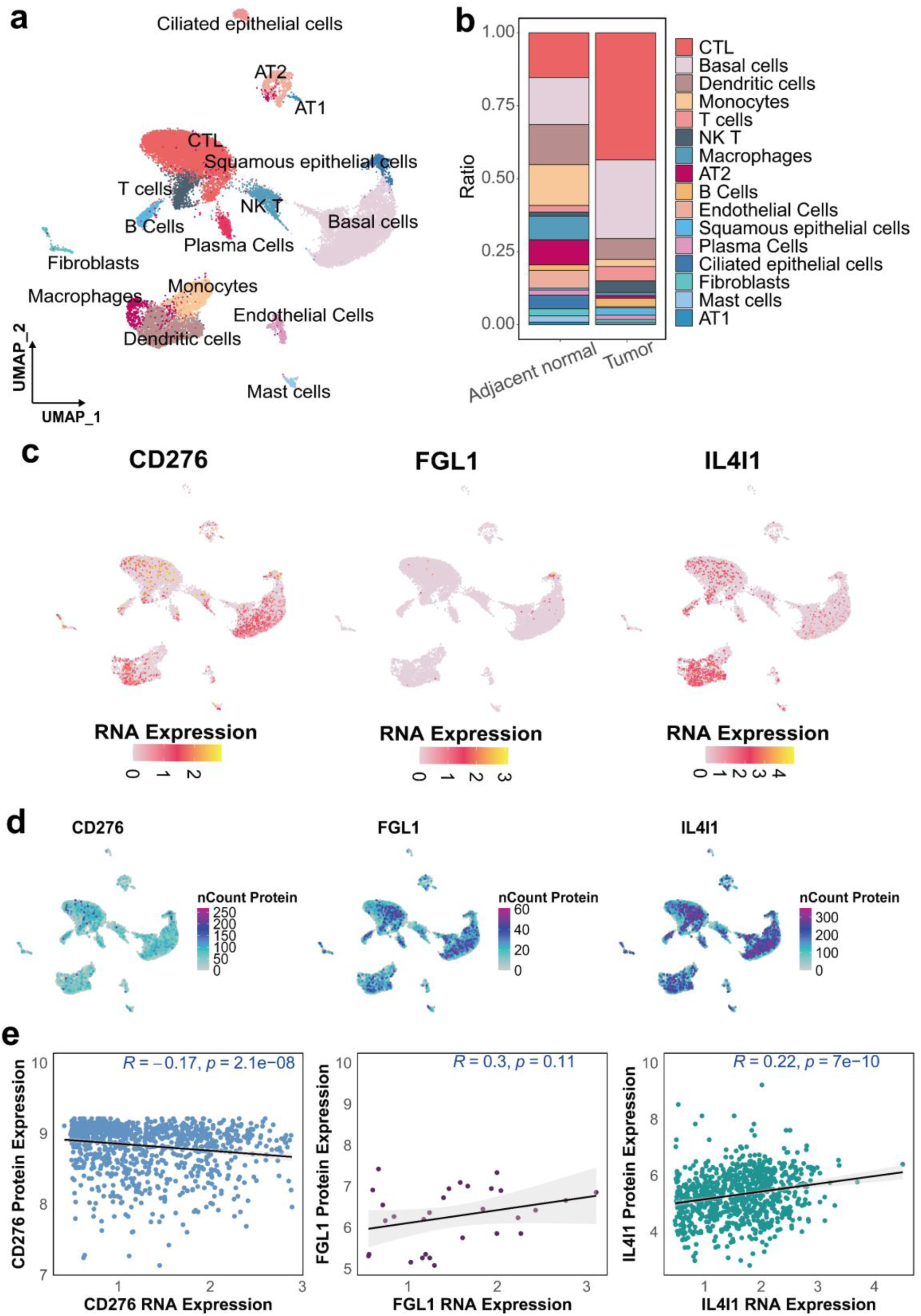
Analysis of cell clustering and expression in LUSC. (a) UMAP visualization of the cell population distribution in LUSC sample. (b) Comparison of various cell populations between adjacent normal tissues and tumor tissues. (c) mRNA expression profiles and (d) marker protein expression profiles. (e) Correlation analysis between mRNA and protein expression levels. (N: adjacent normal tissue, T: tumor tissue)

**Figure S16.**
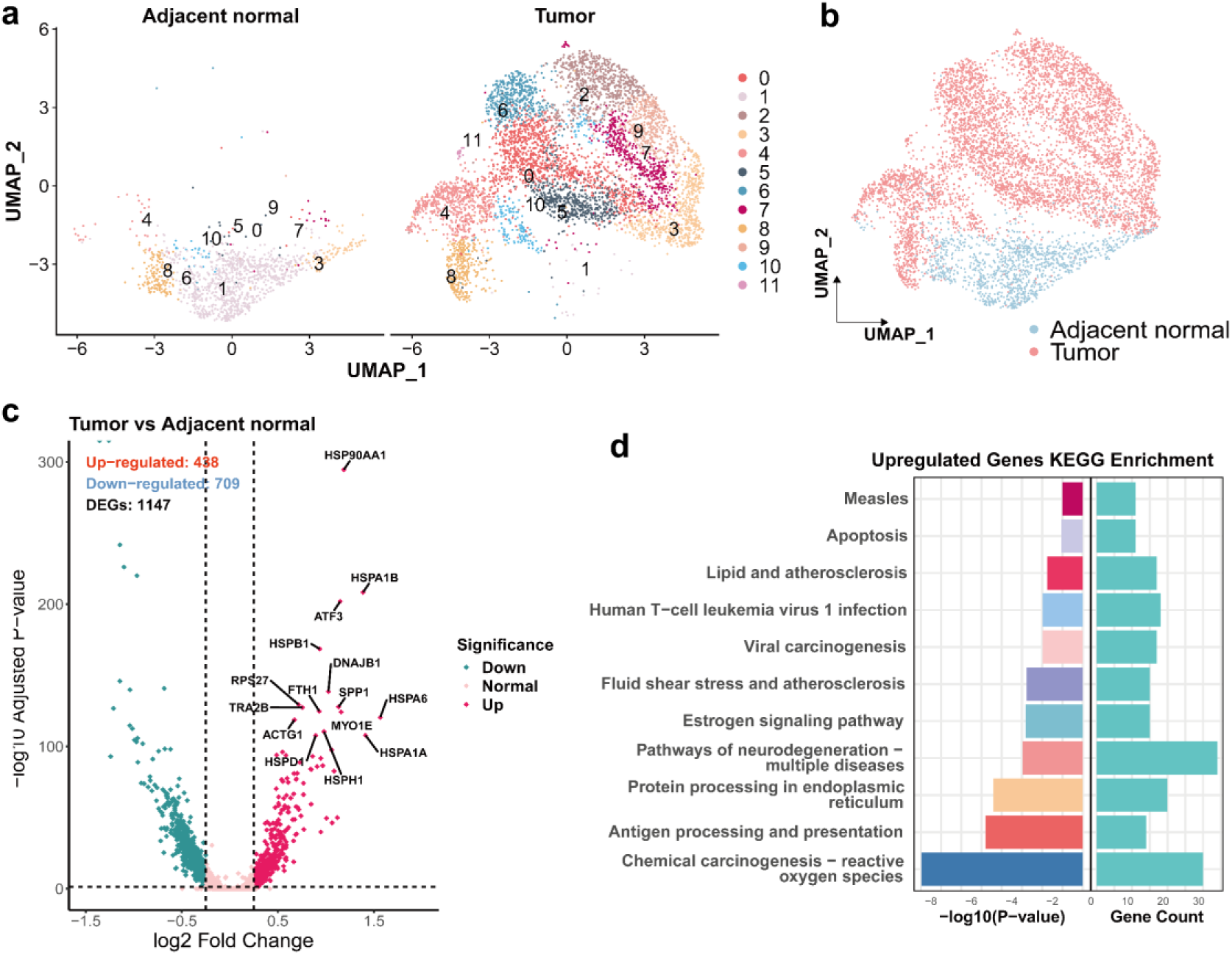
Comparative analysis of basal cells in LUSC tumor and adjacent normal tissues. (a) UMAP visualization comparing the clustering of basal cells in tumor and adjacent normal tissues. (b) Comparison of the distribution of basal cells within the cell population between adjacent normal and tumor tissues. (c) Volcano plot illustrating the DEGs in basal cells between tumor and adjacent normal tissues. (d) KEGG pathway analysis of genes significantly upregulated in tumor-associated basal cells, revealing key pathways associated with tumor progression.

**Figure S17.**
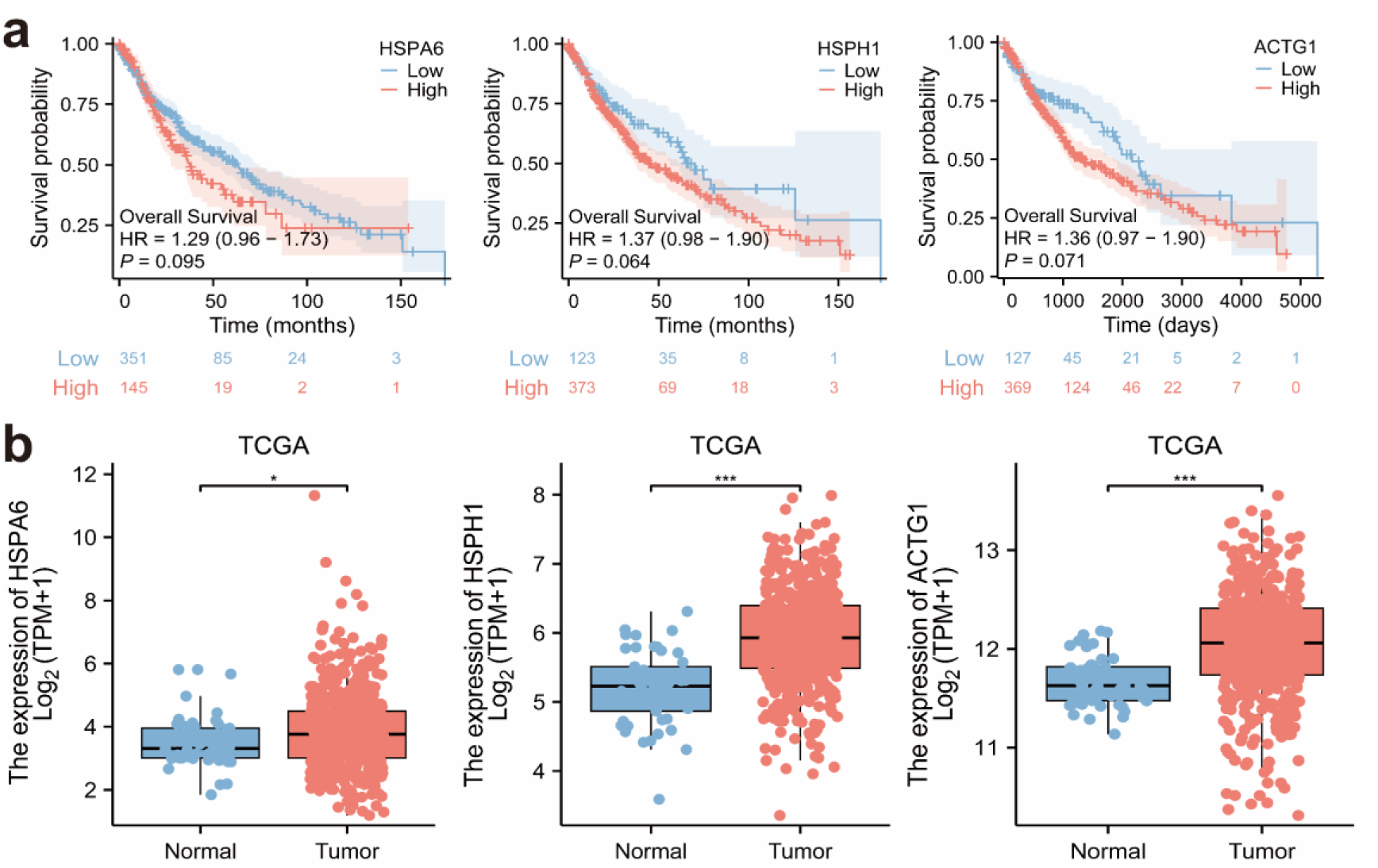
(a) Kaplan‒Meier survival analysis for differentially upregulated genes in basal cells and (b) Analysis of the expression of these genes in adjacent normal and tumor tissues using a TCGA dataset.

