## Supplementary material for "TIP-seq: A Single-cell Multiomics Approach for Simultaneous Transcriptome and Intracellular Protein Profiling": Table S1-S2, Figure S1-S17

Author Information

[1]College of Life Science and Technology, Huazhong University of Science and Technology, Wuhan, 430074, China.

[2]Guangzhou National Laboratory, Guangzhou,510005, China.

[3]College of Life Sciences, University of Chinese Academy of Sciences, Beijing, 101408, China

[4]School of Mechanical & Automotive Engineering, South China University of Technology, Guangzhou, 510641, China

[5]School of Life Sciences, Nankai University, Tianjin, 300071, China

[6]Lead Healthcare (Guangzhou) Co., LTD, Guangzhou, 510005, China

[7]Cancer Hospital & Shenzhen Hospital, Chinese Academy of Medical Sciences and Peking Union Medical College, Shenzhen, 518116, China

[8]Department of Thoracic Surgery, Thoracic Oncology Institute, Peking University People's Hospital, Beijing, 100044, China.

[9] Department of Thoracic Surgery, Shenzhen Second People’s Hospital, Shenzhen, China

[10] Department of Thoracic Surgery, The First Medical Center of Chinese PLA General Hospital, Beijing, 100853, China


[#]These authors contributed equally to this work.

**Table S1:** Sequences of protein barcodes.

| DNA oligo | Sequence (5’-3’) | Modification |
|---|---|---|
| Anti-SCCA-barcode1 | TTGTCTTCCTAAGACCGCTTGGCCTCCGACTT**CGCTCAGTTC**CGATGNNNNNNNNNNGACGCTGCCGACGA | 5’-DBCO-TEG |
| Anti-CD276-barcode2 / Anti-CEA-barcode2 | TTGTCTTCCTAAGACCGCTTGGCCTCCGACTT**TATCTGACCT**CGATGNNNNNNNNNNGACGCTGCCGACGA | 5’-DBCO-TEG |
| Anti-FGL1-barcode3 / Anti-NSE-barcode3 | TTGTCTTCCTAAGACCGCTTGGCCTCCGACTT**TCGGATGTCG**CGATGNNNNNNNNNNGACGCTGCCGACGA | 5’-DBCO-TEG |
| Anti-IL-4I1-barcode4 / Anti-CyFra21-1-barcode4 | TTGTCTTCCTAAGACCGCTTGGCCTCCGACTT**CTTATGGAAT**CGATGNNNNNNNNNNGACGCTGCCGACGA | 5’-DBCO-TEG |
| Anti-ProGRP-barcode5 | TTGTCTTCCTAAGACCGCTTGGCCTCCGACTT**TCCTATTGTG**CGATGNNNNNNNNNNGACGCTGCCGACGA | 5’-DBCO-TEG |
| forward primer (protein) | CGACATGGCTACGATCCGACTT | / |
| reverse primer (protein) | CTTCCTAAGACCGCTTGGCCTC | / |

**Table S2**: Patient information

| | Age | Sex | Histological type | Tumor-Node-Metastasis stage |
|---|---|---|---|---|
| patient 1 | 60 | Man | adenocarcinoma | T1bN0M0, stage IA2 |
| patient 2 | 63 | Man | invasive adenocarcinoma, nonmucinous (lepidic, 50%; acinar, 40%; complex glandular, 10%) | T1aN0M0, stage IA1 |
| patient 3 | 65 | Man | invasive adenocarcinoma, nonmucinous (solid, 70%; complex glandular, 20%; lepidic, 5%; micropapillary, 5%) | T1bN0M0, stage IA2 |
| patient 4 | 54 | Woman | invasive adenocarcinoma, nonmucinous (acinar, 80%; lepidic, 20%) | T1aN0M0, stage IA1 |
| patient 5 | | Woman | adenocarcinoma | IIIA |
| patient 6 | 41 | Man | invasive adenocarcinoma, nonmucinous (acinar, 90%; lepidic, 10%) | T1bN0M0, stage IA2 |
| patient 7 | 71 | Woman | adenocarcinoma (acinar, 50%; solid, 30%; papillary, 10%; micropapillary, 10%) | T1cN0M0, stage IA3 |
| patient 8 | 63 | Woman | invasive adenocarcinoma, nonmucinous (complex glandular, 70%; solid, 50%; acinar, 20%) | T1aN1M0, stage IIA |
| patient 9 | 66 | Man | squamous cell carcinoma | T1bN0M0, stage IA2 |

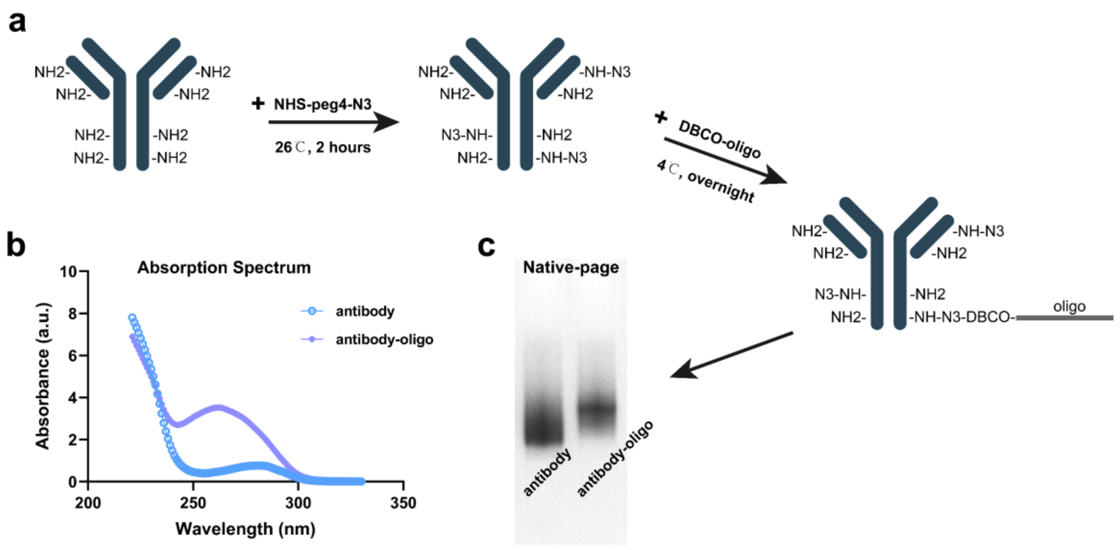


**Figure S1**. (a) A schematic representation illustrating the conjugation process of DBCO-modified oligonucleotides and antibodies. (b) The observed shift in the absorption peak toward 260 nm serves as evidence for the successful conjugation of oligonucleotides to the antibodies. (c) A comparative native gel image depicting the AOCs alongside unmodified antibodies.

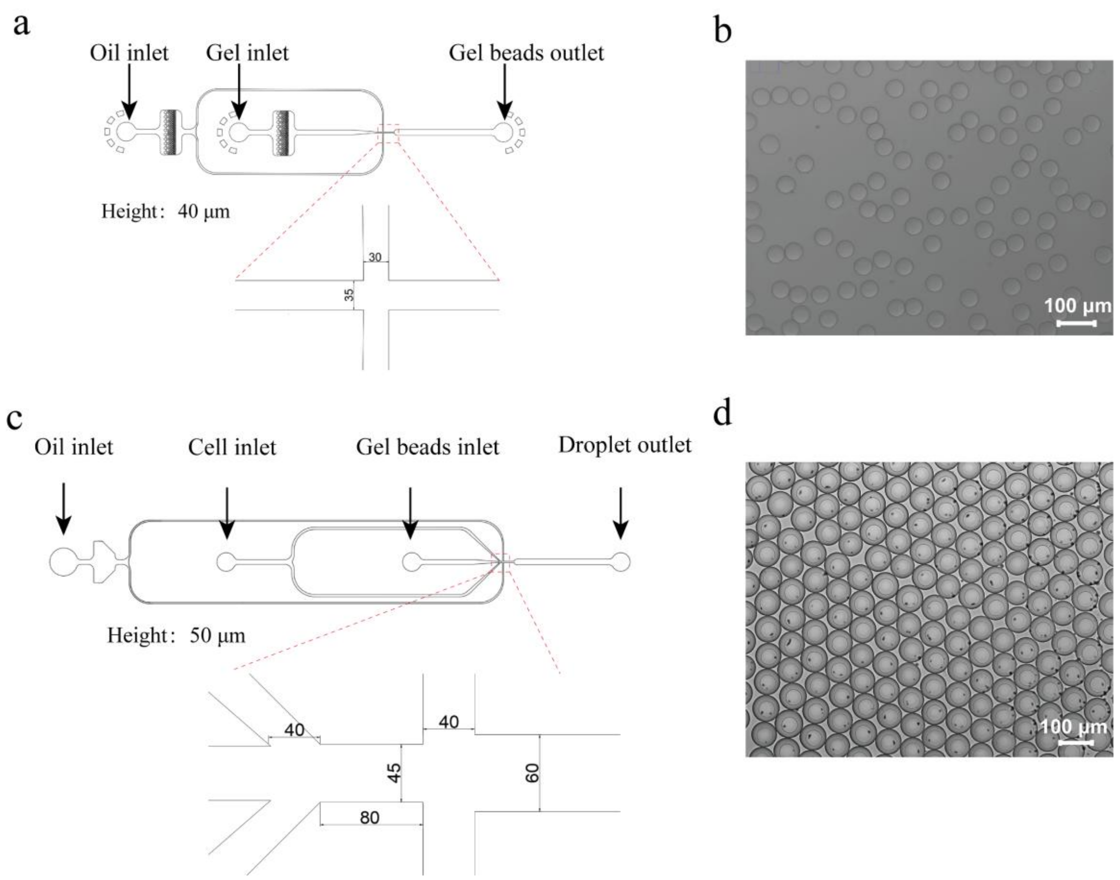


**Figure S2**. (a) Schematic representation of the microfluidic chip designed for hydrogel bead generation. Height: 40 μm. (b) Microscopy image of the hydrogel beads. Scale bar:100 μm. (c) Schematic representation of the microfluidic chip designed for cell encapsulation. Height: 50 μm. (d) Microscopy image of the encapsulated droplets. Scale bar:100 μm.

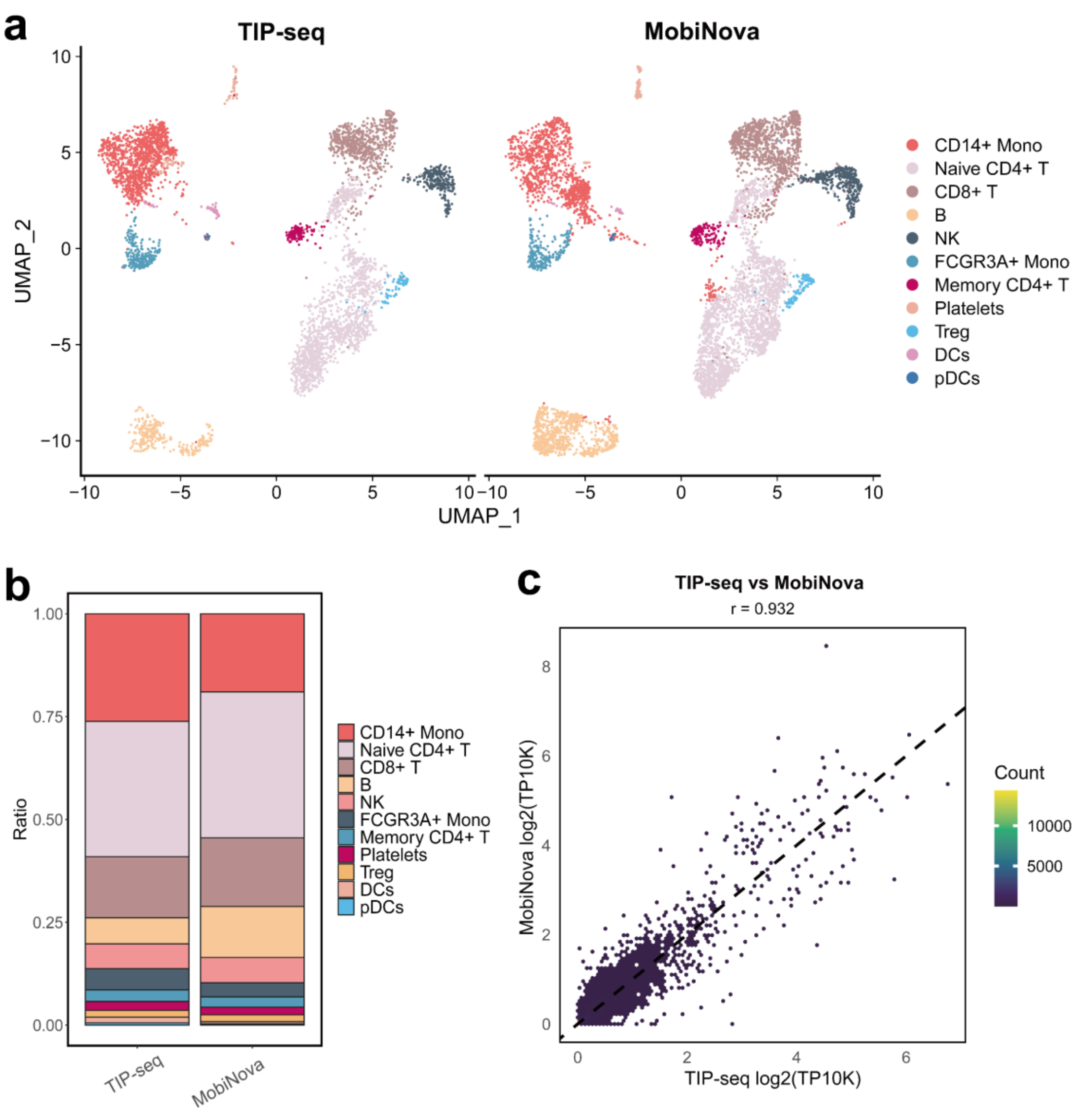


**Figure S3:** Comparative analysis of TIP-seq and MobiNova in PBMC clustering. (a) Comparative evaluation of cell clustering outcomes between TIP-seq and 10x Genomics in PBMC samples. (b) Assessment of the ability of the two technologies to detect cell proportions. (c) Correlation analysis between the two methodologies.

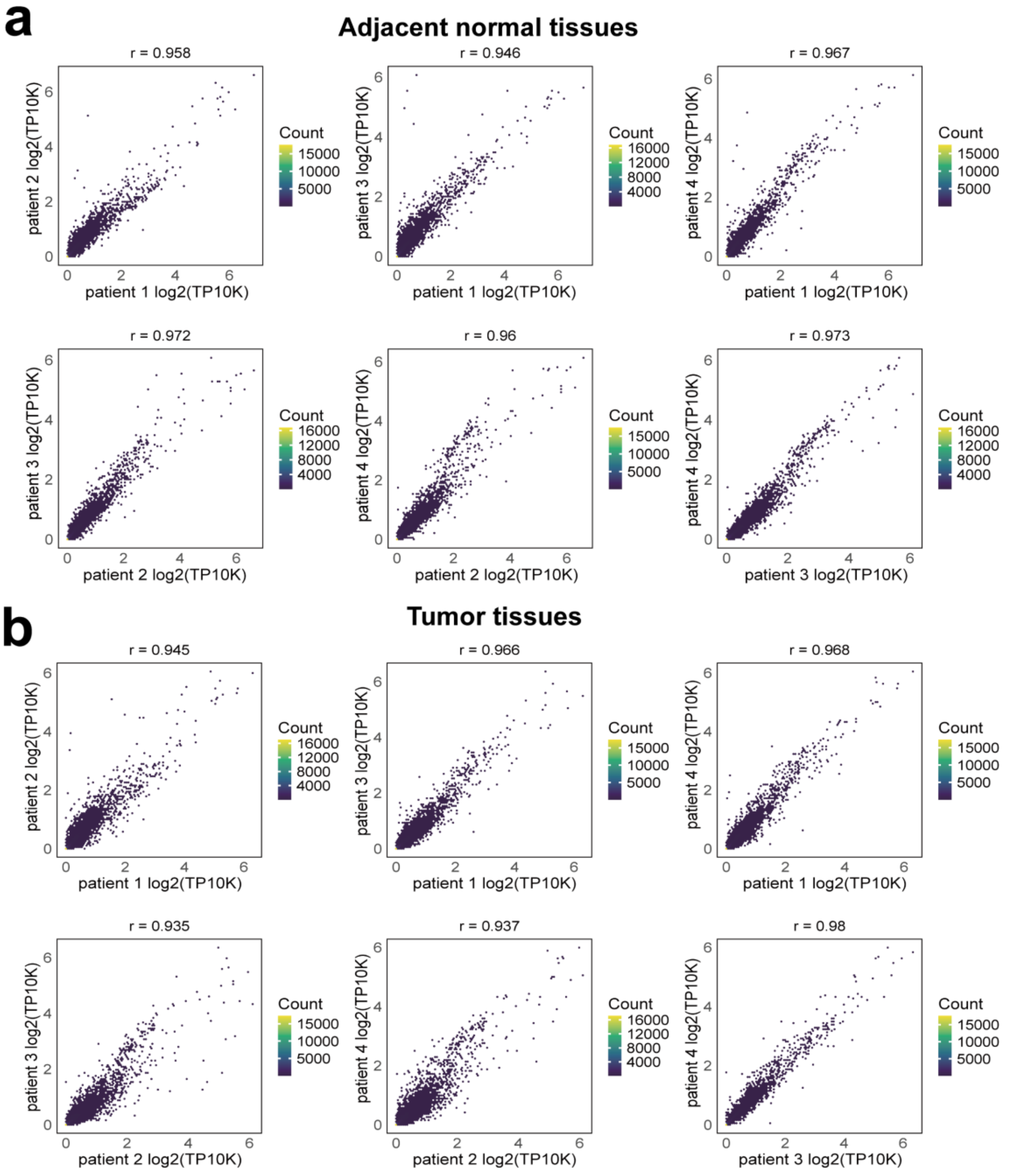


**Figure S4.** Analysis of transcriptomic correlation across NSCLC samples. (a) Correlation analysis of adjacent normal tissues from patients 1-4. (b) Correlation analysis of tumor tissues from patients 1-4.

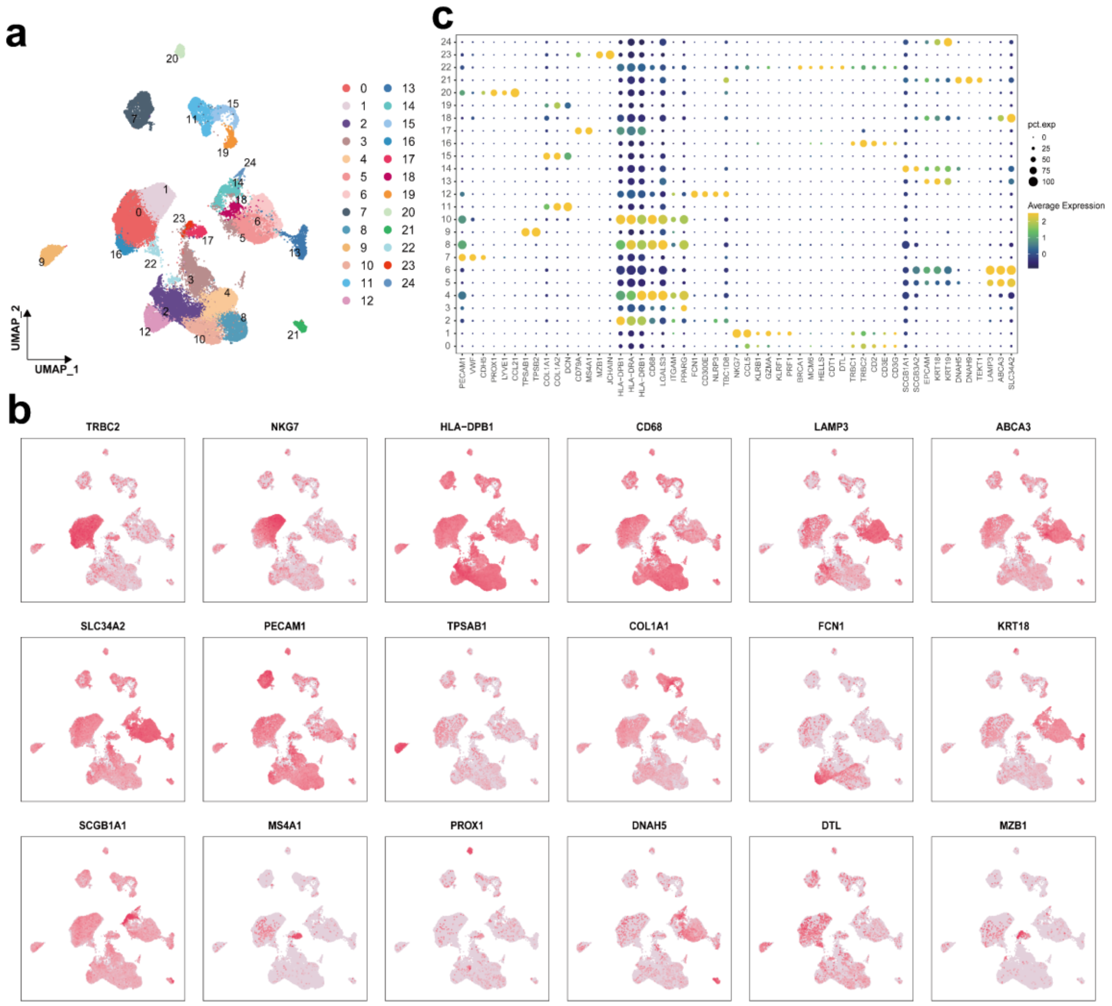


**Figure S5.** Single-cell transcriptomic profiling of NSCLC tissues. (a) UMAP visualization of cell clusters identified in NSCLC tissues. (b) Dot plot displaying the expression levels of key genes across various cell clusters, where the size of each dot represents the percentage of cells expressing the gene and the color intensity reflects the expression level. (c) Expression patterns of marker genes in their respective cell clusters, demonstrating cell-type-specific gene signatures.

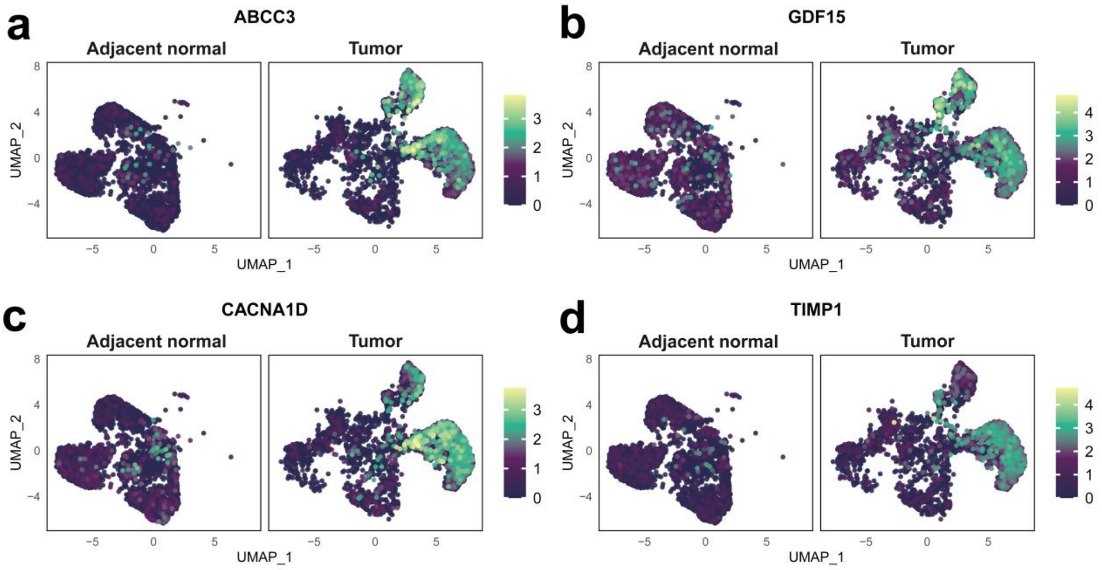


**Figure S6.** Comparative distribution of the four genes whose expression was upregulated in AT2 cells between adjacent normal and tumor tissues. (a) ABCC3: ATP binding cassette subfamily C member 3. (b) GDF15: growth differentiation factor 15. (c) CACNA1D: calcium voltage-gated channel subunit alpha1 D. (d) TIMP1: tissue inhibitor of metal protease 1.

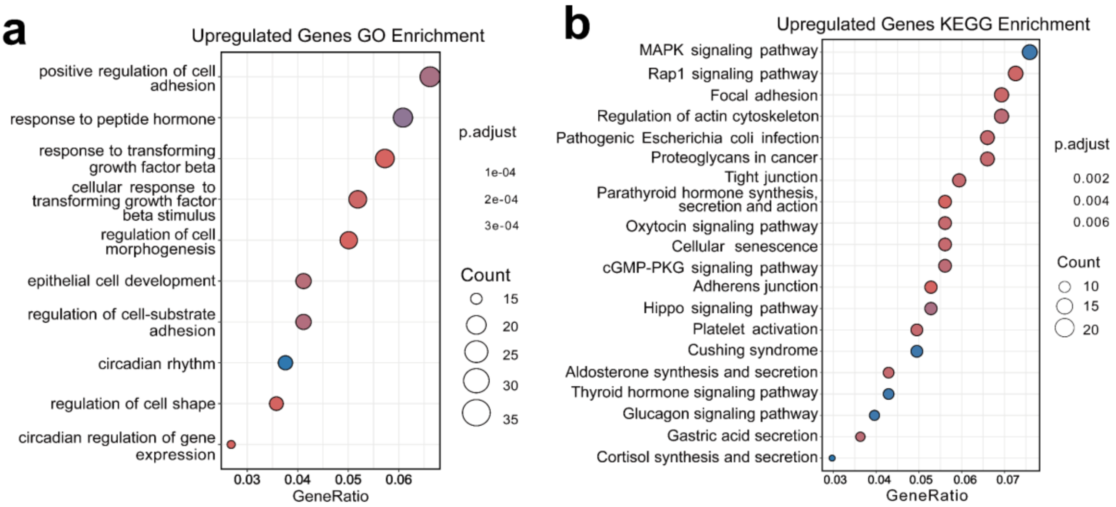


**Figure S7.** (a) GO functional enrichment analysis of biological processes associated with the upregulated genes in AT2 cells. (b) KEGG pathway enrichment analysis of the upregulated genes in AT2 cells.

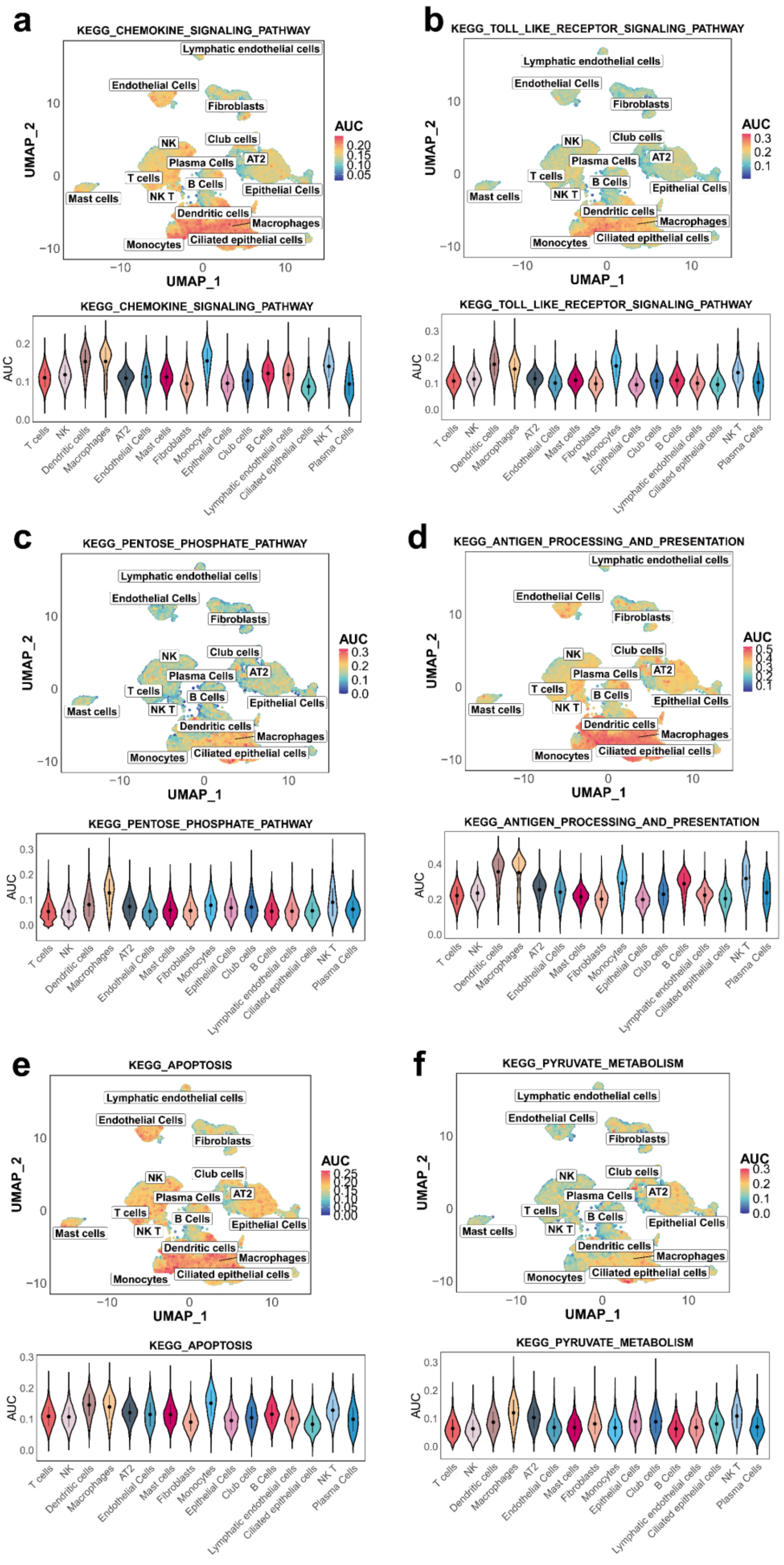
a
KEGG_CHEMOKINE_SIGNALING_PATHWAY
b
KEGG_TOLL_LIKE_RECEPTOR_SIGNALING_PATHWAY
c
KEGG_PENTOSE_PHOSPHATE_PATHWAY
d
KEGG_ANTIGEN_PROCESSING_AND_PRESENTATION
e
KEGG_APOPTOSIS
f
KEGG_PYRUVATE_METABOLISM
Lymphatic endothelial cells
Endothelial Cells
Fibroblasts
NK
Club cells
Plasma Cells
AT2
T cells
B Cells
Mast cells
NK T
Epithelial Cells
Dendritic cells
Macrophages
Monocytes
Ciliated epithelial cells
UMAP_1
UMAP_2
AUC

**Figure S8.** UMAP and violin plots were used to evaluate the expression patterns and diagnostic utility of key signaling pathways across various cell types in tumor and adjacent normal tissues. The violin plots illustrate the area under the curve (AUC) values, which indicate the diagnostic potential of these pathways, with a particular emphasis on macrophages. The pathways analyzed included: (a) chemokine signaling-related pathways, (b) the toll-like receptor signaling pathway, (c) the pentose phosphate pathway, (d) pathways associated with antigen processing and presentation, (e) pathway related to apoptosis, and (f) pyruvate metabolic pathways.

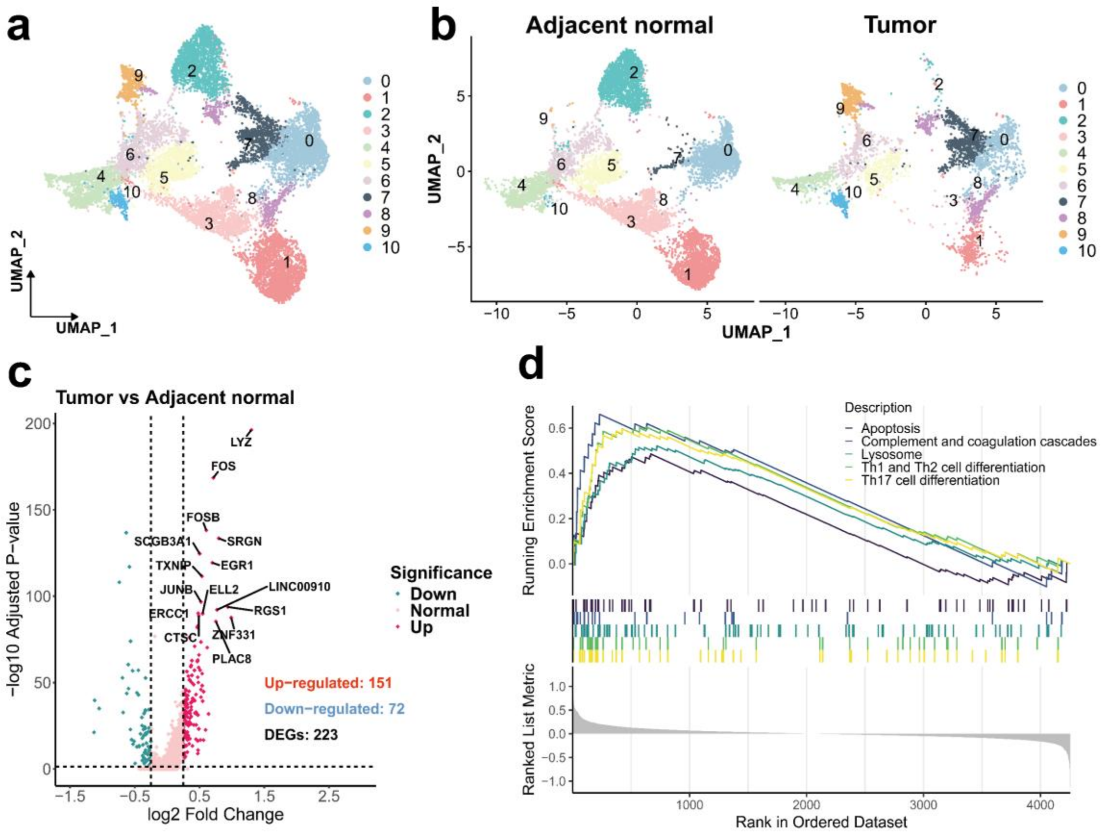


**Figure S9.** Analysis of macrophage traits in LUAD samples. (a) UMAP visualization showing macrophage clusters. (b) Comparative analysis of macrophage distribution in adjacent normal versus tumor tissues. (c) Analysis of gene expression differences in macrophages. (d) Gene set enrichment analysis (GSEA) of the principal signaling pathways activated in macrophages within the TME.

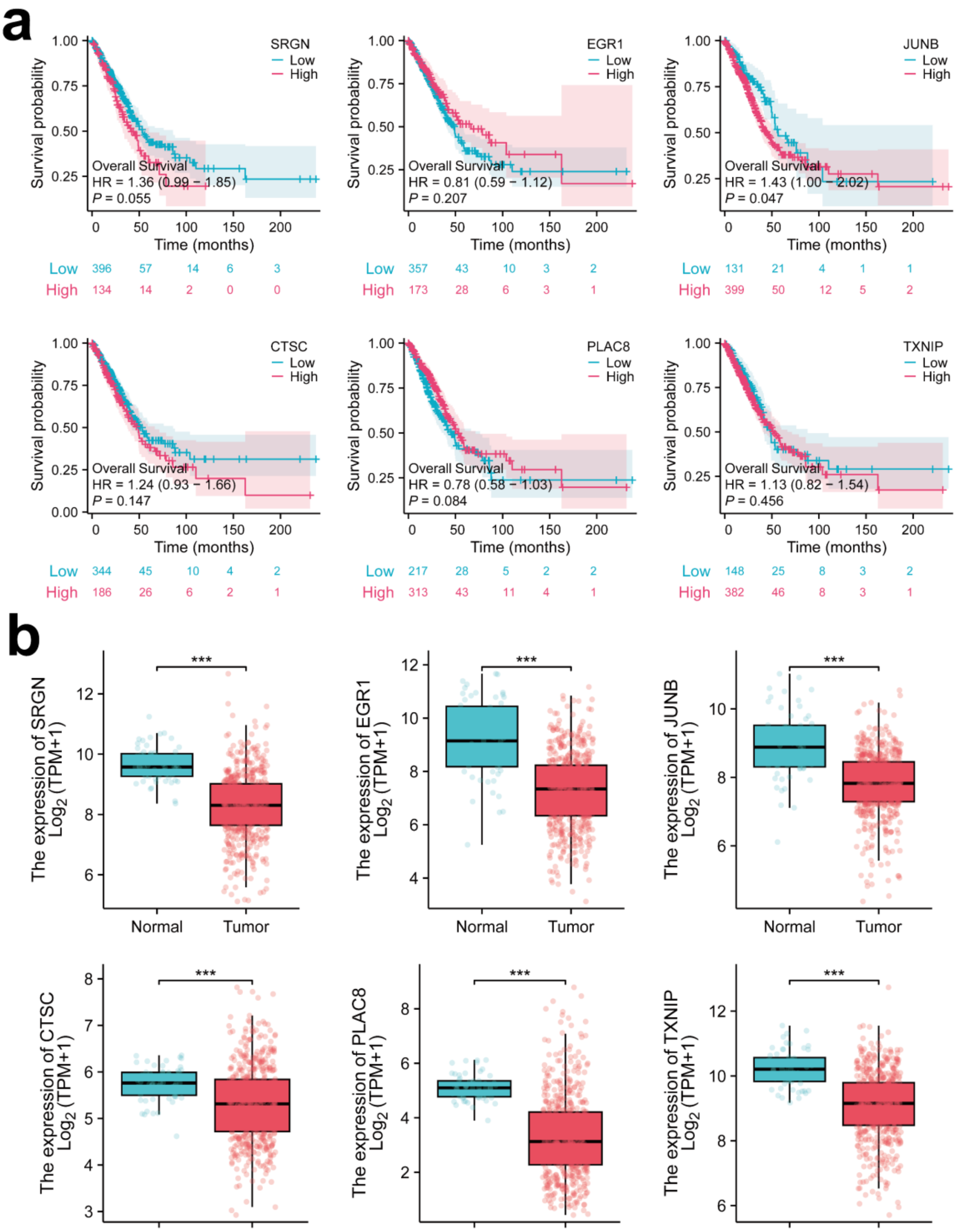


**Figure S10.** (a) Kaplan–Meier survival analysis showing the survival disparities between cohorts with high and low expression of the identified differentially upregulated genes in macrophages. (b) Comparative analysis of genes with elevated expression levels in macrophages on the basis of data derived from the TCGA.

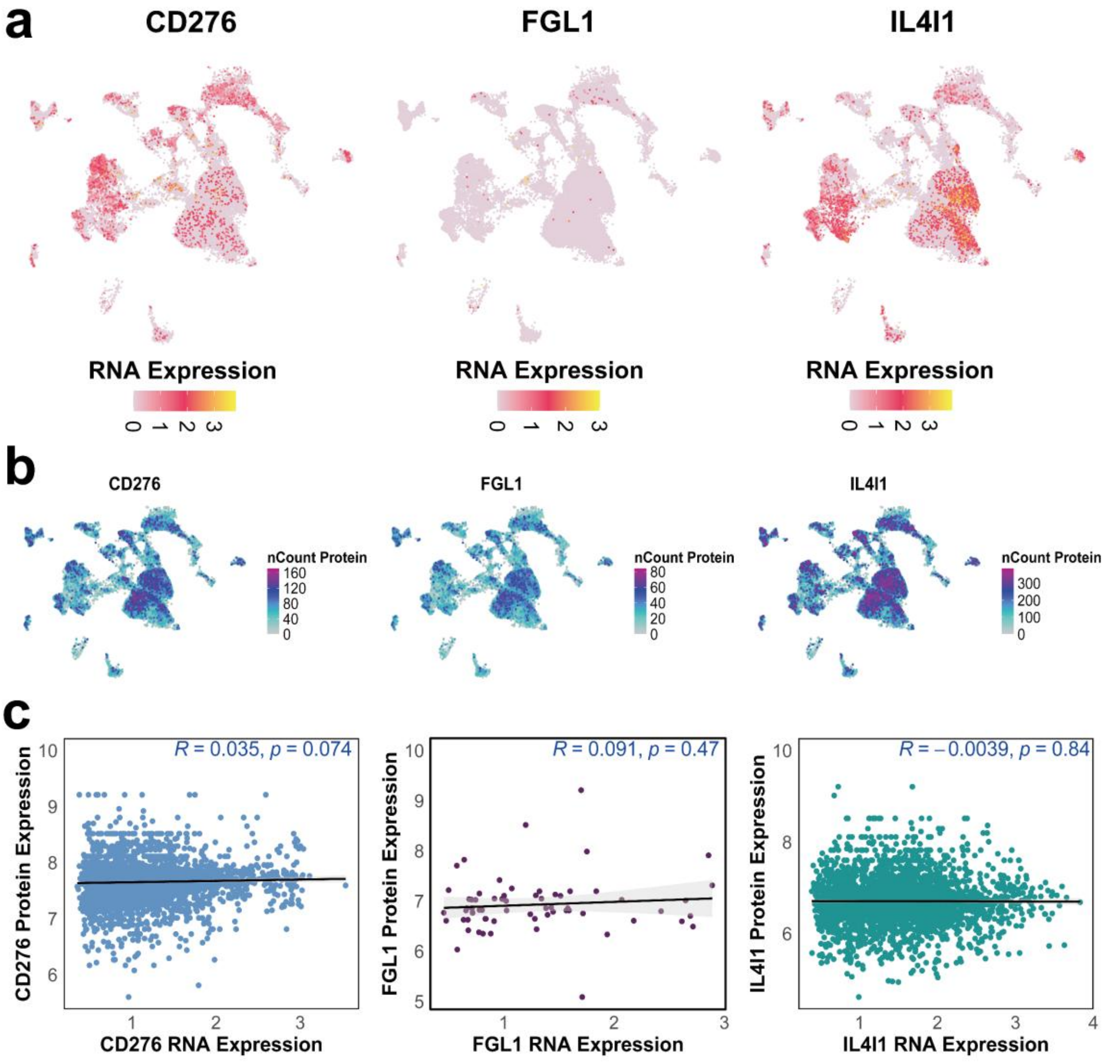


**Figure S11:** (a) Distribution of the RNA expression levels of CD276, FGL1, and IL4I1 in adjacent normal tissues. (b) Corresponding distribution of protein expression levels. (c) Scatter plots depicting the correlation between RNA and protein expression levels for each gene.

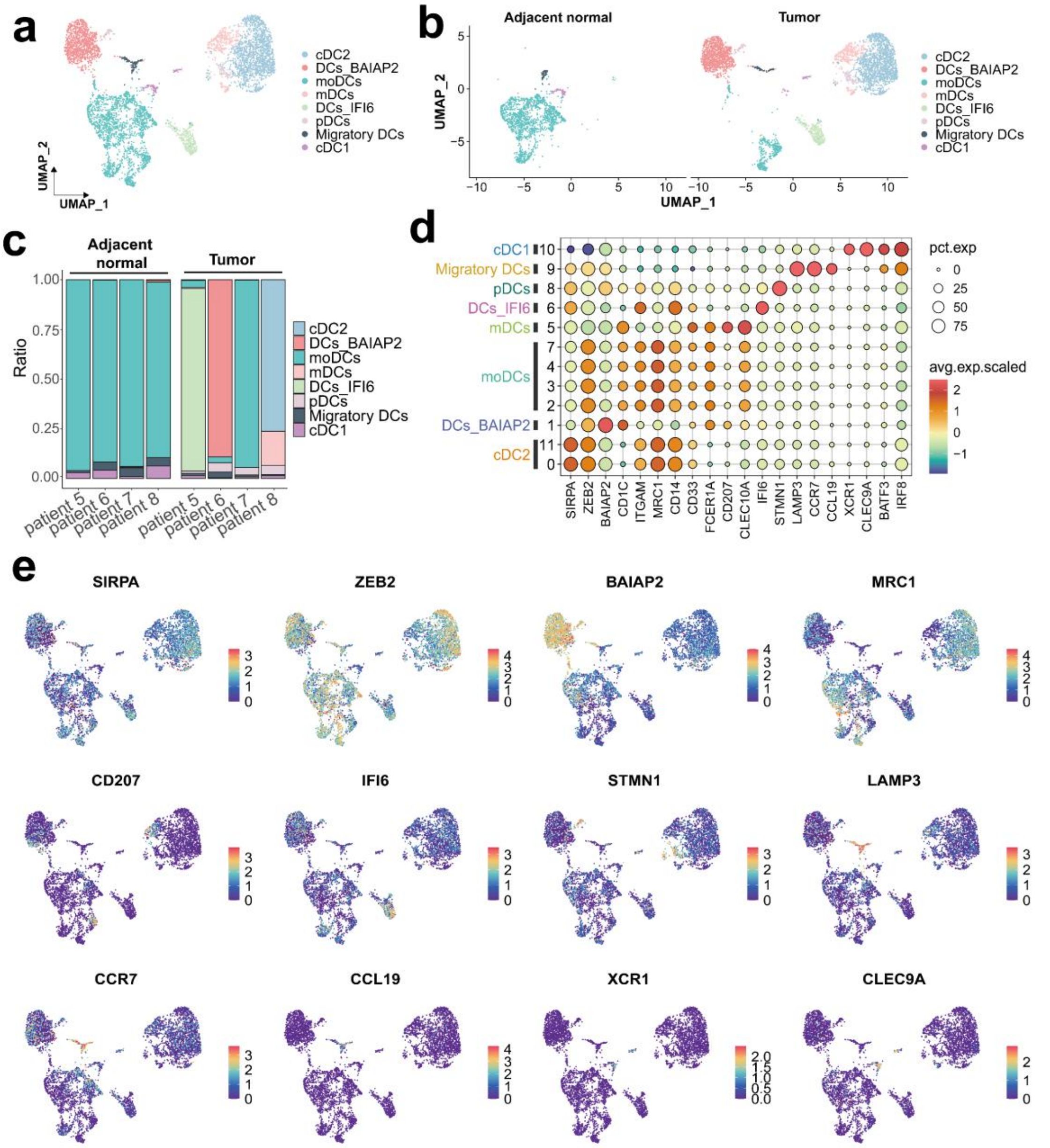


**Figure S12.** Heterogeneity of dendritic cell subpopulations in lung cancer. (a) UMAP visualization of DC subpopulations in lung cancer tissues, revealing distinct clusters and heterogeneity. (b) Comparison of DC subpopulation distributions between normal and lung cancer tissues, showing enrichment of certain subsets in tumors. (c) Proportional analysis of DC subpopulations across different samples, highlighting interindividual heterogeneity in the immune microenvironment. (d) Dot plot displaying distinct expression profiles of DC subpopulation-specific genes, indicating functional diversity. (e) UMAP visualization of the spatial distribution of marker genes in DC subpopulations, highlighting functional diversity within the TME.

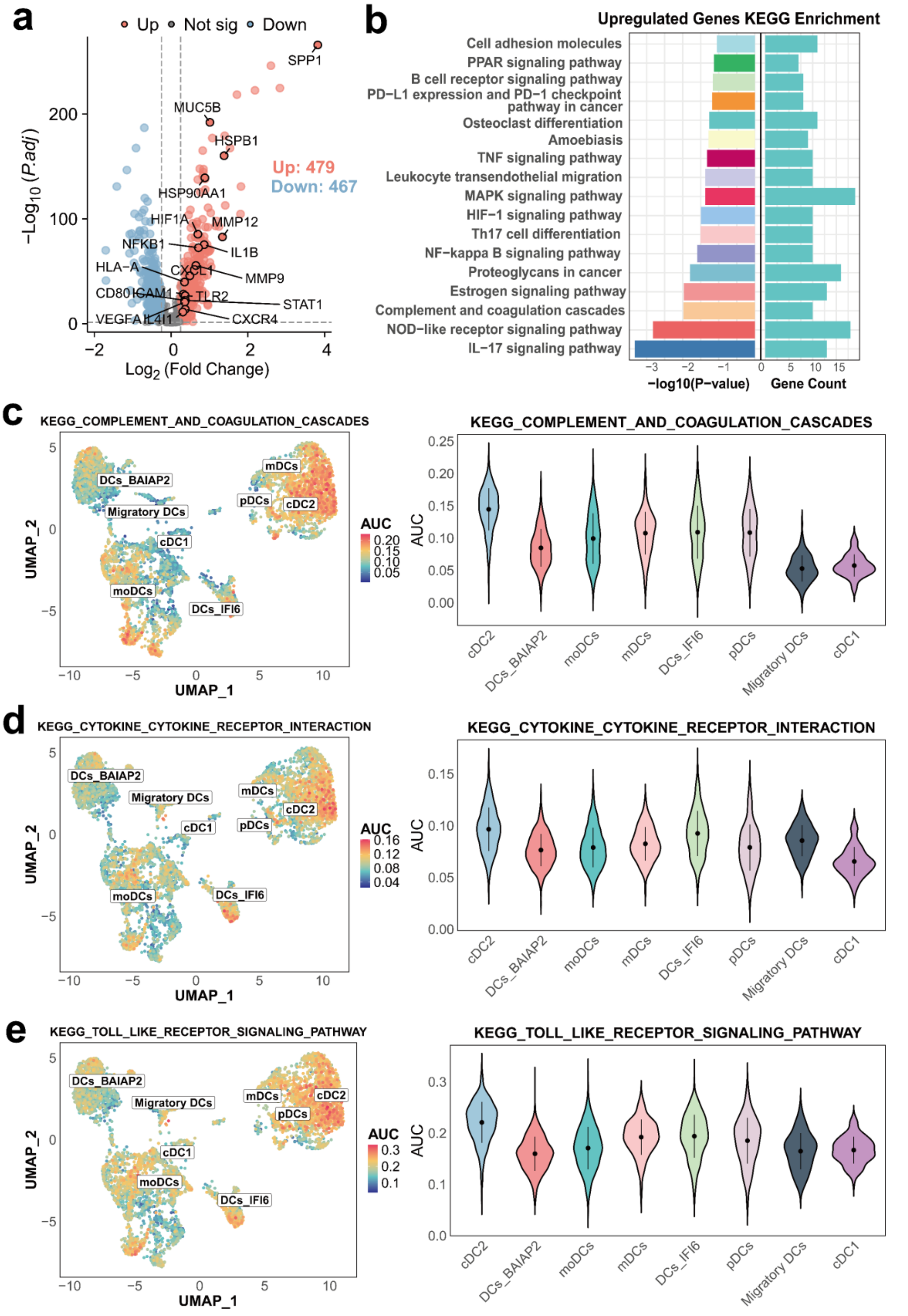


**Figure S13.** Differential expression analysis reveals immune-related signaling pathway activation in lung cancer. (a) Volcano plot showing genes that are differentially expressed between lung cancer and normal tissues, highlighting key upregulated and downregulated genes.

(b) KEGG analysis revealed significant enrichment of immune-related pathways, including pathways associated with complement and coagulation cascades, cytokine–cytokine receptor interaction, and toll–like receptor signaling, in lung cancer tissues. (c) UMAP visualization of complement and coagulation cascade pathway activation in different DC subpopulations, with AUC scores used to quantify pathway activity levels. (d) UMAP visualization of cytokine–cytokine receptor interaction pathway activation in different DC subpopulations, highlighting distinct activation patterns. (e) UMAP visualization of toll–like receptor signaling pathway activation in different DC subpopulations, with AUC scores indicating pathway activity levels.

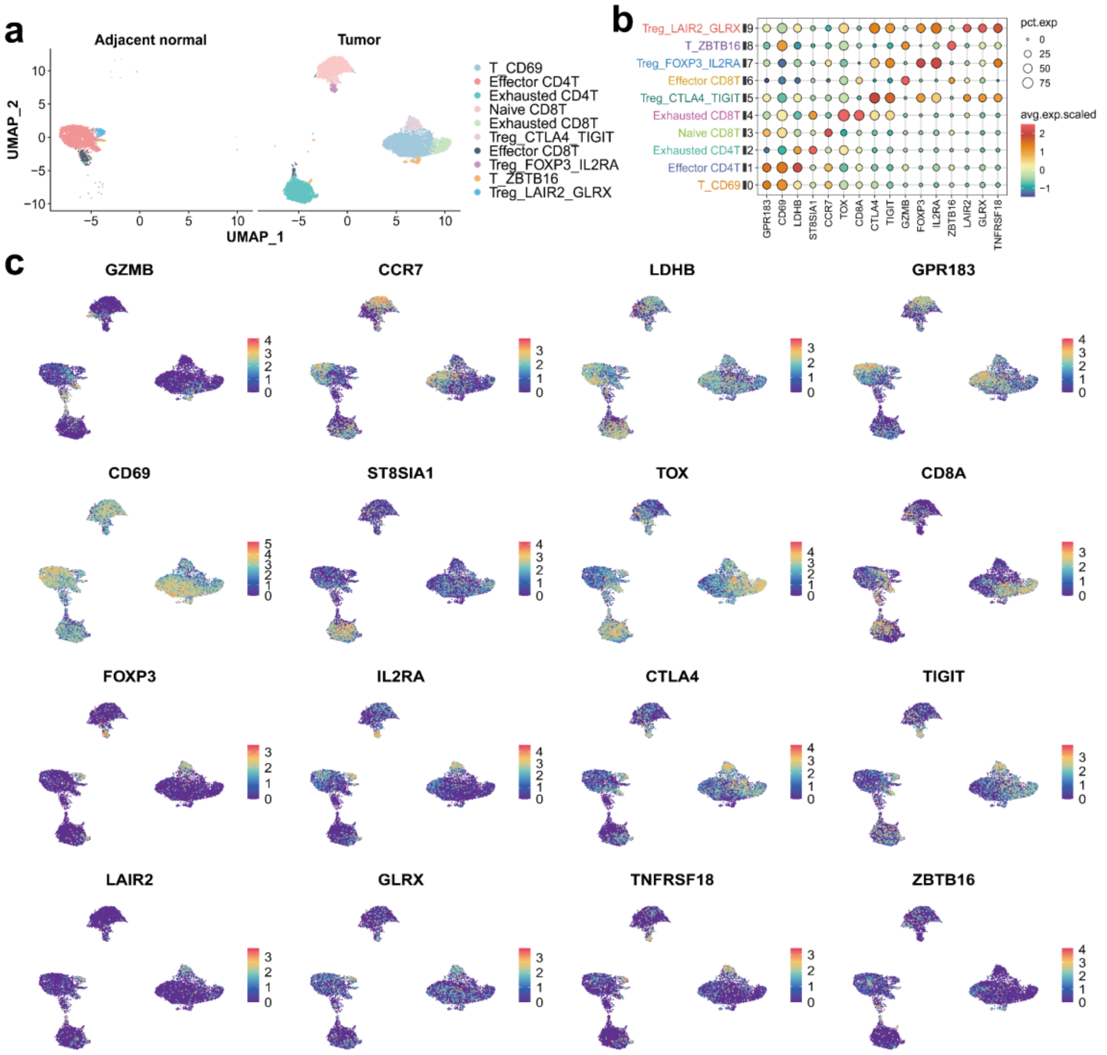


**Figure S14:** T-cell subpopulation heterogeneity and key gene expression in lung cancer. (a) UMAP visualization of T-cell subpopulations in normal and lung cancer tissues, revealing distinct clusters and heterogeneity. (b) Dot plot showing the expression abundance and proportion of functional genes in T-cell subpopulations, highlighting distinct profiles and functional diversity. (c) UMAP visualization of spatial expression patterns of marker genes in T-cell subpopulations, highlighting functional diversity within the TME.

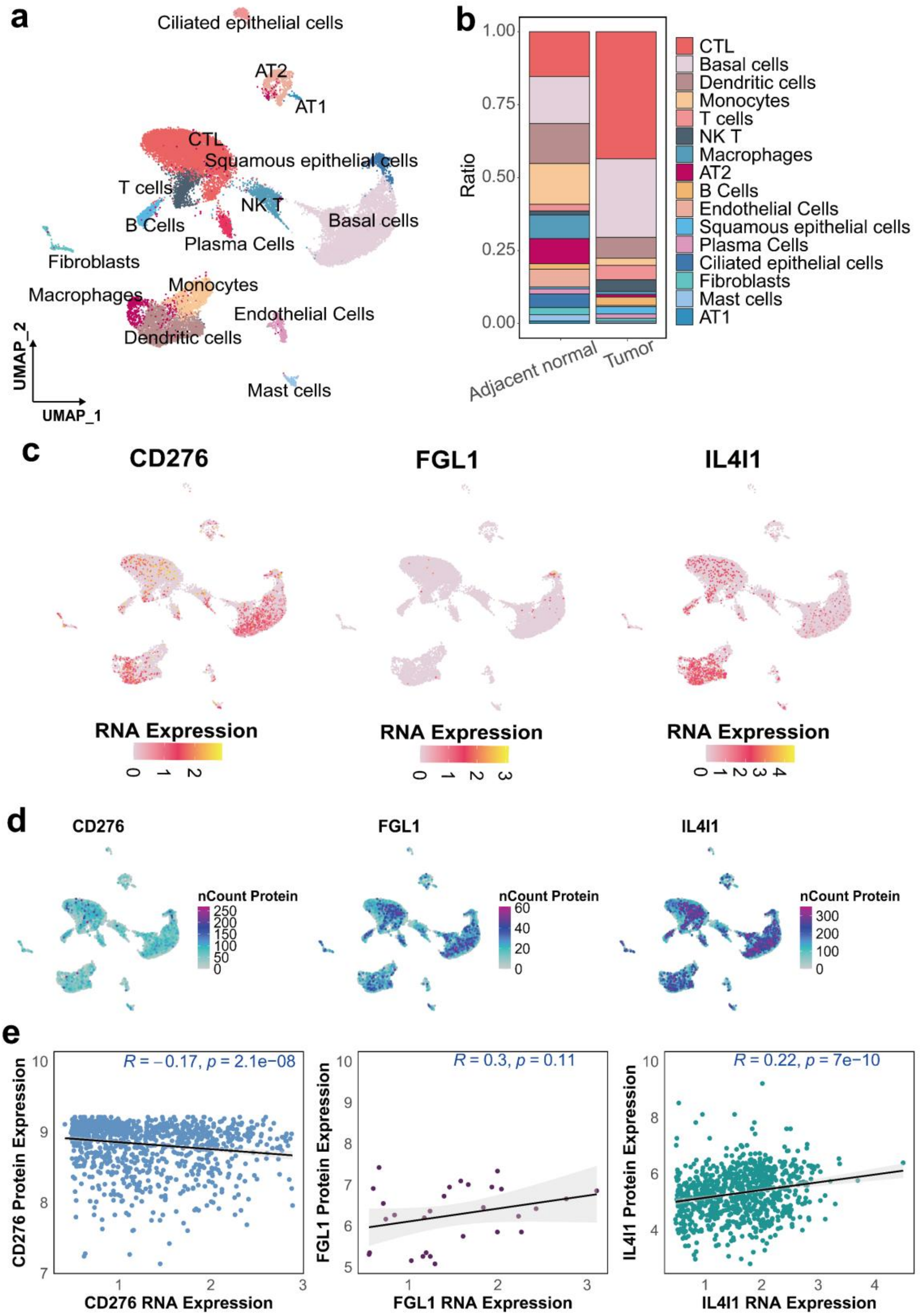

**Figure S15.** Analysis of cell clustering and expression in LUSC. (a) UMAP visualization of the cell population distribution in LUSC sample. (b) Comparison of various cell populations between adjacent normal tissues and tumor tissues. (c) mRNA expression profiles and (d) marker protein expression profiles. (e) Correlation analysis between mRNA and protein expression levels. (N: adjacent normal tissue, T: tumor tissue)

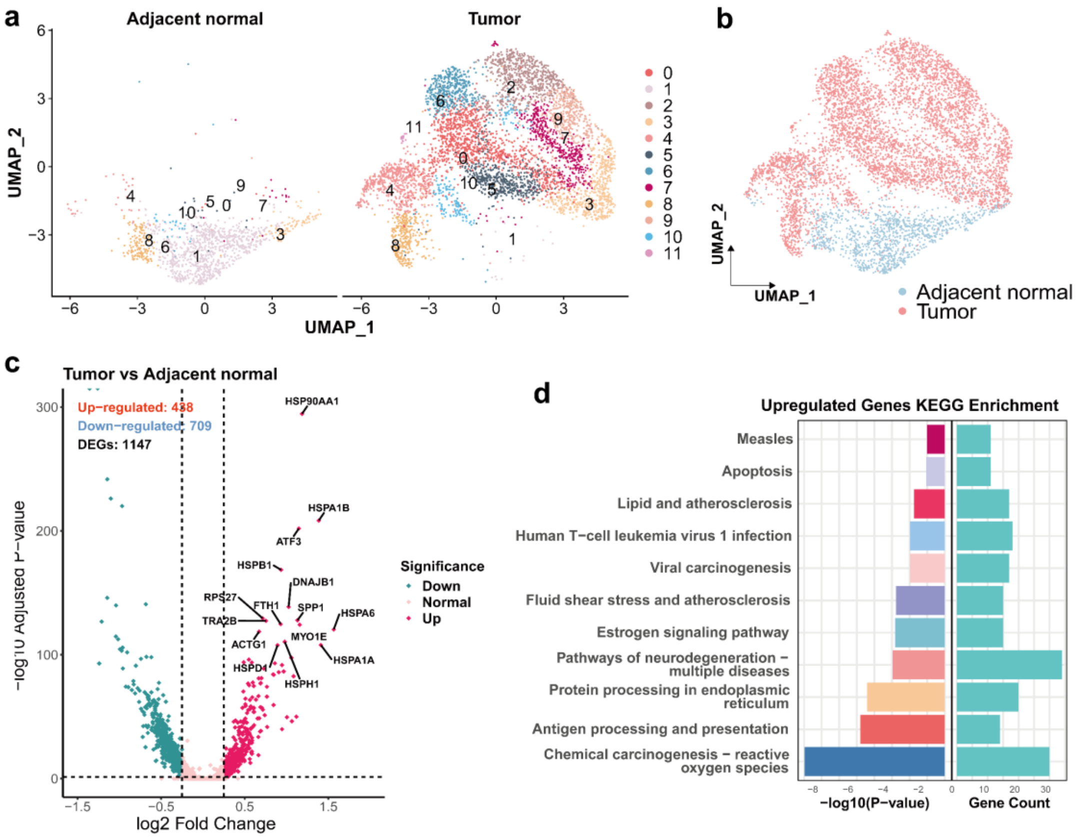


**Figure S16.** Comparative analysis of basal cells in LUSC tumor and adjacent normal tissues. (a) UMAP visualization comparing the clustering of basal cells in tumor and adjacent normal tissues. (b) Comparison of the distribution of basal cells within the cell population between adjacent normal and tumor tissues. (c) Volcano plot illustrating the DEGs in basal cells between tumor and adjacent normal tissues. (d) KEGG pathway analysis of genes significantly upregulated in tumor-associated basal cells, revealing key pathways associated with tumor progression.

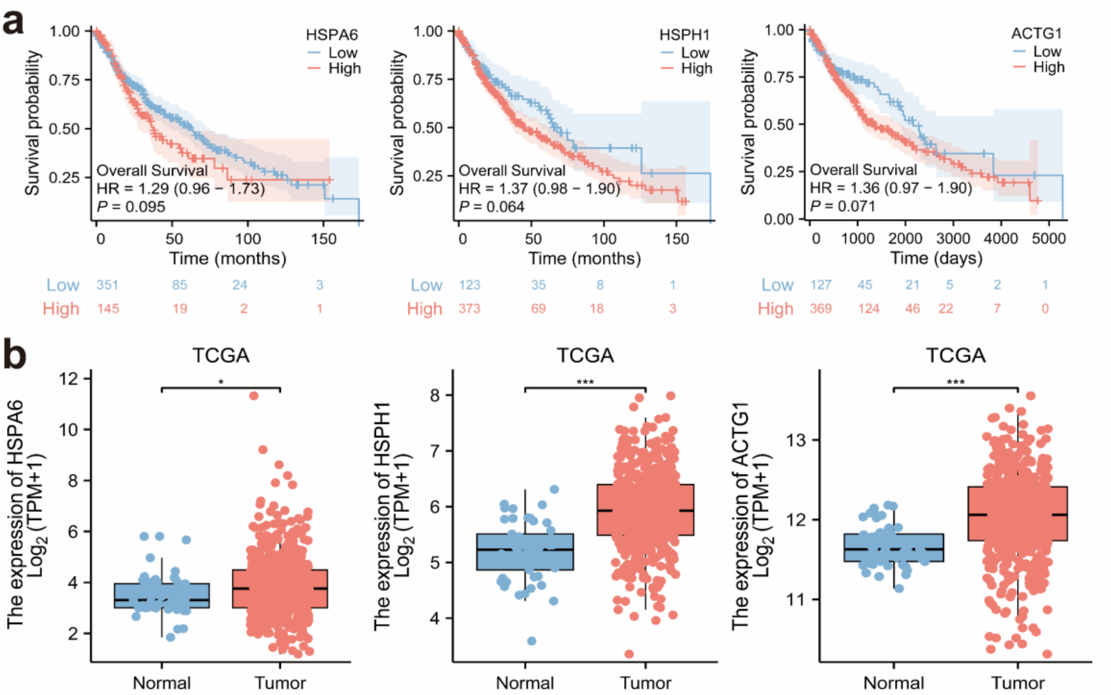


**Figure S17.** (a) Kaplan–Meier survival analysis for differentially upregulated genes in basal cells and (b) Analysis of the expression of these genes in adjacent normal and tumor tissues using a TCGA dataset.
